# Expanding the diversity of bacterioplankton isolates and modeling isolation efficacy with large scale dilution-to-extinction cultivation

**DOI:** 10.1101/2020.04.17.046896

**Authors:** Michael W. Henson, V. Celeste Lanclos, David M. Pitre, Jessica Lee Weckhorst, Anna M. Lucchesi, Chuankai Cheng, Ben Temperton, J. Cameron Thrash

## Abstract

Cultivated bacterioplankton representatives from diverse lineages and locations are essential for microbiology, but the large majority of taxa either remain uncultivated or lack isolates from diverse geographic locales. We paired large scale dilution-to-extinction (DTE) cultivation with microbial community analysis and modeling to expand the phylogenetic and geographic diversity of cultivated bacterioplankton and to evaluate DTE cultivation success. Here, we report results from 17 DTE experiments totaling 7,820 individual incubations over three years, yielding 328 repeatably transferable isolates. Comparison of isolates to microbial community data of source waters indicated that we successfully isolated 5% of the observed bacterioplankton community throughout the study. 43% and 26% of our isolates matched operational taxonomic units and amplicon single nucleotide variants, respectively, within the top 50 most abundant taxa. Isolates included those from previously uncultivated clades such as SAR11 LD12 and *Actinobacteria* acIV, as well as geographically novel members from other ecologically important groups like SAR11 subclade IIIa, SAR116, and others; providing the first isolates in eight putatively new genera and seven putatively new species. Using a newly developed DTE cultivation model, we evaluated taxon viability by comparing relative abundance with cultivation success. The model i) revealed the minimum attempts required for successful isolation of taxa amenable to growth on our media, and ii) identified possible subpopulation viability variation in abundant taxa such as SAR11 that likely impacts cultivation success. By incorporating viability in experimental design, we can now statistically constrain the effort necessary for successful cultivation of specific taxa on a defined medium.

**Importance:** Even before the coining of the term “great plate count anomaly” in the 1980s, scientists had noted the discrepancy between the number of microorganisms observed under the microscope and the number of colonies that grew on traditional agar media. New cultivation approaches have reduced this disparity, resulting in the isolation of some of the “most wanted” bacterial lineages. Nevertheless, the vast majority of microorganisms remain uncultured, hampering progress towards answering fundamental biological questions about many important microorganisms. Furthermore, few studies have evaluated the underlying factors influencing cultivation success, limiting our ability to improve cultivation efficacy. Our work details the use of dilution-to-extinction (DTE) cultivation to expand the phylogenetic and geographic diversity of available axenic cultures. We also provide a new model of the DTE approach that uses cultivation results and natural abundance information to predict taxon-specific viability and iteratively constrain DTE experimental design to improve cultivation success.

## Introduction

Axenic cultures of environmentally important microorganisms are critical for fundamental microbiological investigation, including generating physiological information about environmental tolerances, determining organismal-specific metabolic and growth rates, testing hypotheses generated from *in situ ‘*omics observations, and experimentally examining microbial interactions. Research using important microbial isolates has been critical to a number of discoveries such as defining microorganisms involved in surface ocean methane saturation (1–3), the role of proteorhodopsin in maintaining cellular functions during states of carbon starvation (4, 5), the complete nitrification of ammonia within a single organism (6), and identifying novel metabolites and antibiotics (7, 10). However, the vast majority of taxa remain uncultivated (11–13), restricting valuable experimentation on such topics as genes of unknown function, the role of analogous gene substitutions in overcoming auxotrophy, and the multifaceted interactions occurring in the environment inferred from sequence data (11, 14–16).

The quest to bring new microorganisms into culture, and the recognition that traditional agar-plate based approaches have limited success (17–19), have compelled numerous methodological advances spanning a wide variety of techniques like diffusion chambers, microdroplet encapsulation, and slow acclimatization of cells to artificial media (20–25). Dilution-to-extinction (DTE) cultivation using sterile seawater as the medium has also proven highly successful for isolating bacterioplankton (26–32). Pioneered by Don Button and colleagues for the cultivation of oligotrophic bacteria, this method essentially pre-isolates organisms after serial dilution by separating individual or small groups of cells into their own incubation vessel (32, 33). This prevents slow-growing, obligately oligotrophic bacterioplankton from being outcompeted by faster-growing organisms, as would occur in enrichment-based isolation methods like those that would target aerobic heterotrophs. It is also a practical method for taxa that cannot grow on solid media. Natural seawater media provide these taxa with the same chemical surroundings from which they are collected, reducing the burden of anticipating all the relevant compounds required for growth (33).

Improvements to DTE cultivation in multiple labs have increased the number of inoculated wells and decreased the time needed to detect growth (26, 28, 34), thereby earning the moniker “high-throughput culturing” (26, 28). We (35) and others (30) have also adapted DTE culturing by incorporating artificial media in place of natural seawater media to successfully isolate abundant bacterioplankton. Thus far, DTE culturing has led to isolation of many numerically abundant marine and freshwater groups such as marine SAR11 *Alphaproteobacteria* (28, 29, 34–36), the freshwater SAR11 LD12 clade (29), SUP05/Arctic96BD-19 *Gammaproteobacteria* (37–39), OM43 *Betaproteobacteria* (26, 27, 31, 40, 41), HIMB11-Type Roseobacter spp. (35, 42), numerous so-called Oligotrophic Marine *Gammaproteobacteria* (43), and acI *Actinobacteria* (44).

Despite the success of DTE cultivation, many taxa continue to elude domestication (11–13, 16). Explanations include a lack of required nutrients or growth factors in media (20, 45–49) and biological phenomena such as dormancy and/or phenotypic heterogeneity within populations (47, 48, 50–56). However, there have been few studies empirically examining the factors underlying isolation success in DTE cultivation experiments (34, 57, 58), restricting our ability to determine the relative importance of methodological vs. biological influences on cultivation reliability for any given organism. Moreover, even for those taxa that we have successfully cultivated, in many cases we lack geographically diverse strains, restricting comparisons of the phenotypic and genomic diversity that may influence taxon-specific cultivability.

We undertook a three-year cultivation effort in the coastal northern Gulf of Mexico (nGOM), from which we lack representatives of many common bacterioplankton groups, to provide new model organisms for investigating microbial function, ecology, biogeography, and evolution. Simultaneously, we paired our cultivation efforts with 16S rRNA gene amplicon analyses to compare cultivation results with the microbial communities in the source waters. We have previously reported on the success of our artificial media in obtaining abundant taxa over the course of the first seven experiments from this campaign (35). Here, we expand our report to include cultivation results from a total of seventeen experiments, and update the classic viability calculations of Button *et al.* (33) with a new model to estimate the viability of individual taxa using relative abundance information. New isolates belonged to cultivated groups in eight putatively novel genera and seven putatively novel species in previously cultivated genera and expanded cultured geographic representation for many important clades like SAR11. Additionally, using model-based predictions, we identified possible taxon-specific viability variation that can influence cultivation success. By incorporating these new viability estimates into the model, our method facilitates statistically-informed experimental design for targeting individual taxa, thereby reducing uncertainty for future culturing work (59).

## Material and Methods

### Sampling

Surface water samples were collected at six different sites once a year for three years, except for Terrebonne Bay, which was collected twice. The sites sampled were Lake Borgne (LKB; Shell Beach, LA), Bay Pomme d’Or (JLB; Buras, LA), Terrebonne Bay (TBON; Cocodrie, LA), Atchafalaya River Delta (ARD; Franklin, LA), Freshwater City (FWC; Kaplan, LA), and Calcasieu Jetties (CJ; Cameron, LA) (lat/long coordinates provided in Table S1). Water collection for biogeochemical and biological analysis followed the protocol in (35). Briefly, we collected surface water in a sterile, acid-washed polycarbonate bottle. Duplicate 120 ml water samples were filtered serially through 2.7 μm Whatman GF/D (GE Healthcare, Little Chalfort, UK) and 0.22 μm Sterivex (Millipore, Massachusetts, USA) filters and placed on ice until transferred to −20°C in the laboratory (maximum 3 hours on ice). The University of Washington Marine Chemistry Laboratory analyzed duplicate subsamples of 50 ml 0.22 μm-filtered water collected in sterile 50 ml falcon tubes (VWR, Pennsylvania, USA) for concentrations of SiO_4_, PO_4_^3−^, NH_4_^+^, NO_3_^−^, and NO_2_^−^. Samples for cell counts were filtered through a 2.7 μm GF/D filter, fixed with 10% formaldehyde, and stored on ice until enumeration (maximum 3 hours). Temperature, salinity, pH, and dissolved oxygen were measured using a handheld YSI 556 multiprobe system (YSI Inc., Ohio, USA). All metadata is available in Table S1.

### Dilution-to-extinction culturing and propagation

Isolation, propagation, and identification of isolates were completed as previously reported (29, 35, 60). A subsample of 2.7 μm filtered surface water was stained with 1X SYBR Green (Lonza, Basal, Switzerland) using a repeat pipettor and disposal tip (Gilson, Wisconsin, USA), and enumerated using a Guava Easycyte 5HT HPL flow cytometer (Millipore, Massachusetts, USA) as described (60). After serial dilution to a predicted 1-3 cells·μl^−1^, 2 μl water was inoculated into five, 2 mL 96-well PTFE plates (Radleys, Essex, UK) containing 1.7 ml artificial seawater medium (Table S1) using a 20 uL multichannel pipet (Gilson, Wisconsin, USA) to achieve an estimated 1-3 cells·well^−1^ (Table 1). The salinity of the medium was chosen to match *in situ* salinity after experiment JLB (January 2015) (Tables 1, S1). After year two, a second generation of media, designated MWH, was designed to incorporate additional important osmolytes, reduced sulfur compounds, and other constituents (Tables 1, S1) potentially necessary for *in vitro* growth of uncultivated clades (49, 61–67). The four corner wells of each plate were left uninoculated as negative controls for every experiment. Plates were covered using sterile, PTFE-coated silicon 96-well plate mats (Thermo Scientific, Massachusetts, USA). Cultures were incubated at *in situ* temperatures (Table S1) in the dark for three to six weeks and evaluated for positive growth (> 10^4^ cells·ml^−1^) by flow cytometry. 200 μl from positive wells was transferred using a 200 μL single channel pipet (Gilson, Wisconsin, USA) to duplicate 125 ml polycarbonate flasks (Corning, New York, USA) containing 50 ml of medium (29, 35, 60). At FWC, FWC2, JLB2c, and JLB3, not all positive wells were transferred because of the large number of positive wells. At each site, 48/301, 60/403, 60/103, and 60/146 of the positive wells were transferred, respectively, selected using flow cytometry signatures with < 10^2^ green fluorescence and < 10^2^ side scatter that maximized our chances of isolating small microorganisms that encompass many of the most abundant and most wanted taxa, like SAR11, using our settings (60).

**Table 1.** Cultivation statistics, including whole community viability estimates

| Site | Date | *n* | *z* | *p* | $\lambda$ | *V* * (ASE) | CV | our model:<br>estimated #<br>wells with 1<br>cell<br>(bootstrapped<br>median: (xx-xx)<br>95% CI) if<br>*V*=1 ** | our<br>model:<br>*V*<sub>est</sub>: min-max 95%<br>CI *** | *in situ*<br>salinity | Medium<br>salinity | Medium | Study |
| --- | --- | --- | --- | --- | --- | --- | --- | --- | --- | --- | --- | --- | --- |
| CJ | Sep 2014 | 460 | 15 | 0.033 | 1.27 | 2.6 (0.67) | 0.259 | 164 (144-185) | 1.5-4.2 | 24.6 | 34.8 | JWAMPF<br>e | (35) |
| ARD | Nov 2014 | 460 | 1 | 0.002 | 1.5 | 0.1 (0.15) | 1.451 | 154 (134-174) | 0.1-0.7 | 1.72 | 34.8 | JW1 | (35) |
| JLB | Jan 2015 | 460 | 61 | 0.133 | 1.96 | 7.3 (0.93) | 0.127 | 127 (109-146) | 5.6-9.2 | 26.0 | 34.8 | JW1 | (35) |
| FWC<sup>†</sup> | Mar 2015 | 460 | 301 | 0.654 | 2 | 53.1 (3.2) | 0.06 | 125 (106-143) | 47.1-59.7 | 5.39 | 5.79 | JW4 | (35) |
| LKB | June 2015 | 460 | 15 | 0.033 | 1.8 | 1.8 (0.48) | 0.266 | 137 (118-156) | 1.1-3.0 | 2.87 | 5.79 | JW4 | (35) |
| Tbon2 | Aug 2015 | 460 | 41 | 0.089 | 1.56 | 6.0 (0.93) | 0.156 | 151 (132-171) | 4.3-8.1 | 14.2 | 11.6 | JW3 | (35) |
| CJ2 | Oct 2015 | 460 | 61 | 0.133 | 2 | 7.1 (0.91) | 0.128 | 125 (106-143) | 5.6-9.1 | 22.2 | 23.2 | JW2 | (35) |
| FWC2<sup>†</sup> | Apr 2016 | 460 | 403 | 0.876 | 2 | 104.4 (6.2) | 0.059 | 125 (106-143) | >92.3 | 20.9 | 23.2 | JW2 | This study |
| ARD2c | Jun 2016 | 460 | 7 | 0.015 | 2 | 0.8 (0.29) | 0.362 | 125 (106-143) | 0.3-1.5 | 0.18 | 1.45 | JW5 | This study |
| JLB2c<sup>†</sup> | May 2016 | 460 | 103 | 0.224 | 2 | 12.7 (1.25) | 0.099 | 125 (106-143) | 10.3-15.4 | 6.89 | 5.79 | JW4 | This study |
| LKB2 | Jul 2016 | 460 | 39 | 0.085 | 2 | 4.4 (0.71) | 0.161 | 125 (106-143) | 3.2-6.0 | 2.39 | 1.45 | JW5 | This study |
| Tbon3 | Jul 2016 | 460 | 78 | 0.17 | 2 | 9.3 (1.05) | 0.113 | 125 (106-143) | 7.4-11.5 | 17.7 | 34.8 | MWH1 | This study |
| CJ3 | Sep 2016 | 460 | 69 | 0.15 | 2 | 8.1 (0.98) | 0.121 | 125 (106-143) | 6.4-10.2 | 23.7 | 23.2 | MWH2 | This study |
| FWC3 | Nov 2016 | 460 | 27 | 0.059 | 2 | 3.0 (0.58) | 0.194 | 125 (106-143) | 2.0-4.4 | 18.0 | 23.2 | MWH2 | This study |
| ARD3 | Dec 2016 | 460 | 58 | 0.126 | 2 | 6.7 (0.89) | 0.132 | 125 (106-143) | 5.2-8.6 | 3.72 | 1.45 | MWH5 | This study |
| JLB3<sup>†</sup> | Jan 2017 | 460 | 146 | 0.317 | 2 | 19.1 (1.59) | 0.083 | 125 (106-143) | 16.1-22.4 | 12.4 | 11.6 | MWH3 | This study |
| LKB3 | Feb 2017 | 460 | 38 | 0.083 | 2 | 4.3 (0.70) | 0.163 | 125 (106-143) | 3.1-5.8 | 3.55 | 1.45 | MWH5 | This study |

\*Viability according to equation 1, below. Asymptotic Standard Error (ASE) is presented in parentheses.
\*\*Based on 9,999 bootstraps.
\*\*\*Based on 9,999 bootstraps tested at viability increments of 0.1%.
<sup>†</sup>Experiments where a subset of positive wells were transferred.
FWC2 shows the advantage of our method over equation 1 for extreme values.

### Culture identification

Cultures reaching ≥ 1 × 10^5^ cells·ml^−1^ had 35 ml of the 50 ml volume filtered for identification via 16S rRNA gene PCR onto 25 mm 0.22-μm polycarbonate filters (Millipore, Massachusetts, USA). DNA extractions were performed using the MoBio PowerWater DNA kit (QIAGEN, Massachusetts, USA) following the manufacturer’s instructions and eluted in sterile water. The 16S rRNA gene was amplified as previously reported in Henson et al. 2016 (35) and sequenced at Michigan State University Research Technology Support Facility Genomics Core. Evaluation of Sanger sequence quality was performed with 4Peaks (v. 1.7.1) (http://nucleobytes.com/4peaks/) and forward and reverse complement sequences (converted via http://www.bioinformatics.org/sms/rev_comp.html) were assembled where overlap was sufficient using the CAP3 web server (http://doua.prabi.fr/software/cap3).

### Community iTag sequencing, operational taxonomic units, and single nucleotide variants

Sequentially filtered (2.7μm, 0.22μm) duplicate samples were extracted and analyzed using our previously reported protocols and settings (35, 68). We sequenced the 2.7-0.22 μm fraction for this study because this fraction corresponded with the < 2.7 μm communities that were used for the DTE experiments. To avoid batch sequencing effects, DNA from the first seven collections reported in (35) was resequenced with the additional samples from this study (FWC2 and after-Table 1). We targeted the 16S rRNA gene V4 region with the 515F, 806RB primer set (that corrects for poor amplification of taxa like SAR11) (69, 70) using Illumina MiSeq 2 × 250bp paired-end sequencing at Argonne National Laboratories, resulting in 2,343,106 raw reads for the 2.7-0.22 μm fraction. Using Mothur v1.33.3 (71), we clustered 16S rRNA gene amplicons into distinctive OTUs with a 0.03 dissimilarity threshold (OTU_0.03_) and classified them according to the Silva v119 database (72, 73). After these steps, 55,256 distinct OTUs_0.03_ remained. We also used minimum entropy decomposition (MED) to partition reads into fine-scale amplicon single nucleotide variants (ASVs) (74). Reads were first analyzed using Mothur as described above up to the *screen.seqs()* command. The cleaned reads fasta file was converted to MED-compatible headers with the ‘mothur2oligo’ tool renamer.pl from the functions in MicrobeMiseq (https://github.com/DenefLab/MicrobeMiseq) (75) using the fasta output from *screen.seqs()* and the Mothur group file. These curated reads were analyzed using MED (v. 2.1) with the flags −M 60, and -d 1. MED resulted in 2,813 refined ASVs. ASVs were classified in Mothur using *classify.seqs()*, the Silva v119 database, and a cutoff bootstrap value of 80% (76). After classification, we removed ASVs identified as “chloroplast”, “mitochondria”, or “unknown” from the dataset.

### Community analyses

OTU (OTU_0.03_) and ASV abundances were analyzed within the R statistical environment v.3.2.1 following previously published protocols (29, 35, 68). Using the package PhyloSeq (78), OTUs and ASVs were rarefied using the command *rarefy_even_depth()* and OTUs/ASVs without at least two reads in four of the 34 samples (2 sites; ~11%) were removed. This latter cutoff was used to remove potentially spurious OTUs/ASVs resulting from sequencing errors. Our modified PhyloSeq scripts are available on our GitHub repository https://github.com/thrash-lab/Modified-Phyloseq. After filtering, the datasets contained 777 unique OTUs and 1,323 unique ASVs (Table S1). For site-specific community comparisons, beta diversity between sites was examined using Bray-Curtis distances via ordination with non-metric multidimensional scaling (NMDS) (Table S1). The nutrient data were normalized using the R function *scale* which subtracts the mean and divides by the standard deviation for each column. The influence of the transformed environmental parameters on beta diversity was calculated in R with the *envfit* function. Relative abundances of an OTU or ASV from each sample were calculated as the number of reads over the sum of all the reads in that sample. The relative abundance was then averaged between biological duplicates for a given OTU or ASV. To determine the best matching OTU or ASV for a given LSUCC isolate, the OTU representative fasta file, provided by Mothur using *get.oturep*(), and the ASV fasta file were used to create a BLAST database (*makeblastdb*) against which the LSUCC isolate 16S rRNA genes could be searched via blastn (BLAST v 2.2.26) (“OTU_ASVrep_db” - Available as Supplemental Information at https://doi.org/10.6084/m9.figshare.12142098). We designated a LSUCC isolate 16S rRNA gene match with an OTU or ASV sequence based on ≥ 97% or ≥ 99% sequence identity, respectively, as well as a ≥ 247 bp alignment.

### 16S rRNA gene phylogeny

Taxa in the *Alpha*-, *Beta*-, and *Gammaproteobacteria* phylogenies from (35) served as the backbone for the trees in the current work. For places in these trees with poor representation near isolate sequences, additional taxa were selected by searching the 16S rRNA genes of LSUCC isolates against the NCBI nt database online with BLASTn (79) and selecting a variable number of best hits. The *Bacteroidetes* and *Actinobacteria* trees were composed entirely of non-redundant top 100-300 MegaBLAST hits to a local version of the NCBI nt database, accessed August 2018. Sequences were aligned with MUSCLE v3.6 (80) using default settings, culled with TrimAl v1.4.rev22 (81) using the -automated1 flag, and the final alignment was inferred with IQ-TREE v1.6.11 (82) with default settings and −bb 1000 for ultrafast bootstrapping (83). Tips were edited with the nw_rename script within Newick Utilities v1.6 (84) and trees were visualized with Archaeopteryx (85). Fasta files for these trees and the naming keys are available as Supplemental Information at https://doi.org/10.6084/m9.figshare.12142098.

### Assessment of isolate novelty

We quantified taxonomic novelty using BLASTn of our isolate 16S rRNA genes to those of other known isolates collected in three databases: 1) The NCBI nt database (accessed August 2018) - “NCBIdb”; 2) a custom database comprised of sequences from DTE experiments in other labs - “DTEdb”; and 3) a database containing all of our isolate 16S rRNA genes - “LSUCCdb”. The DTEdb and LSUCCdb fasta files are available as Supplemental Information at https://doi.org/10.6084/m9.figshare.12142098. We compared our isolate sequences to these databases as follows:

1. All representative sequences were searched against the nt database using BLASTn (BLAST+v. 2.7.1) with the flags -perc_identity 84, -evalue 1E-6, -task blastn, -outfmt “6 qseqid sseqid pident length slen qlen mismatch evalue bitscore sscinames sblastnames stitle”, and -negative_gilist to remove uncultured and environmental sequences. The negative GI list was obtained by searching `“environmental samples”[organism] OR metagenomes[orgn]”` in the NCBI Nucleotide database (accessed September 12^th^, 2018) and hits were downloaded in GI list format. This negative GI list is available as Supplemental Information at https://doi.org/10.6084/m9.figshare.12142098. The resultant hits from the NCBIdb search were further manually curated to remove sequences classified as single cell genomes, clones, duplicates, and previously deposited LSUCC isolates.
2. We observed that many known HTCC, IMCC, and HIMB isolates that had previously been described as matching our clades (Figs. S1-5) were missing from the resultant lists of nt hits, so we extracted isolate accession numbers from numerous DTE experiments (26–28, 31, 34, 37, 44, 86, 87) from the nt database via *blastdbcmd* and generated a separate DTEdb using *makeblastdb*. Duplicate accession numbers found in the NCBIdb were removed. The same BLASTn settings as in 1) were used to search our isolate sequences against DTEdb. Any match that fell below the lowest percent identity hit to the NCBIdb was removed from the DTEdb search since the match would not have been present in the first NCBIdb search.
3. Finally, using the same BLASTn settings, we compared all pairwise identities of our 328 LSUCC isolate 16S rRNA gene sequences via the LSUCCdb.

The output from these searches is available in Table S1 under the “taxonomic novelty” tab. We placed our LSUCC isolates into 55 taxonomic groups based on sharing ≥ 94% identity and/or their occurrence in monophyletic groups within our 16S rRNA gene trees (Figs. S1-5, see above). For visualization purposes, in groups with multiple isolates we used our chronologically first cultivated isolate as the representative sequence for blastn searches, and these are the top point (100% identity to itself) in each group column of Figure 1. Sequences from the other DTE culture collections were labeled with the corresponding collection name, while all other hits were labeled as “Other”.

Geographic novelty was assessed by manually screening the accession numbers from hits to LSUCC isolates with ≥ 99% 16S rRNA gene sequence identity for the latitude and longitude from a connected publication or location name (e.g. source, country, site) in the NCBI description. LSUCC isolates in the *Janibacter* sp., *Micrococcus* sp., *Altererythrobacter* sp., *Pseudomonas* sp., and *Phycicoccus* sp. groups (16 total isolates) were not assessed because of missing isolation source information and no traceable publication. Isolation locations were plotted for a subset of important taxa (Table S1 “Map_cultivars” tab) using the “LSUCC_cultivar_map.R” available at our GitHub repository https://github.com/thrash-lab/Cultivar-novelty-map.

**Figure 1.**
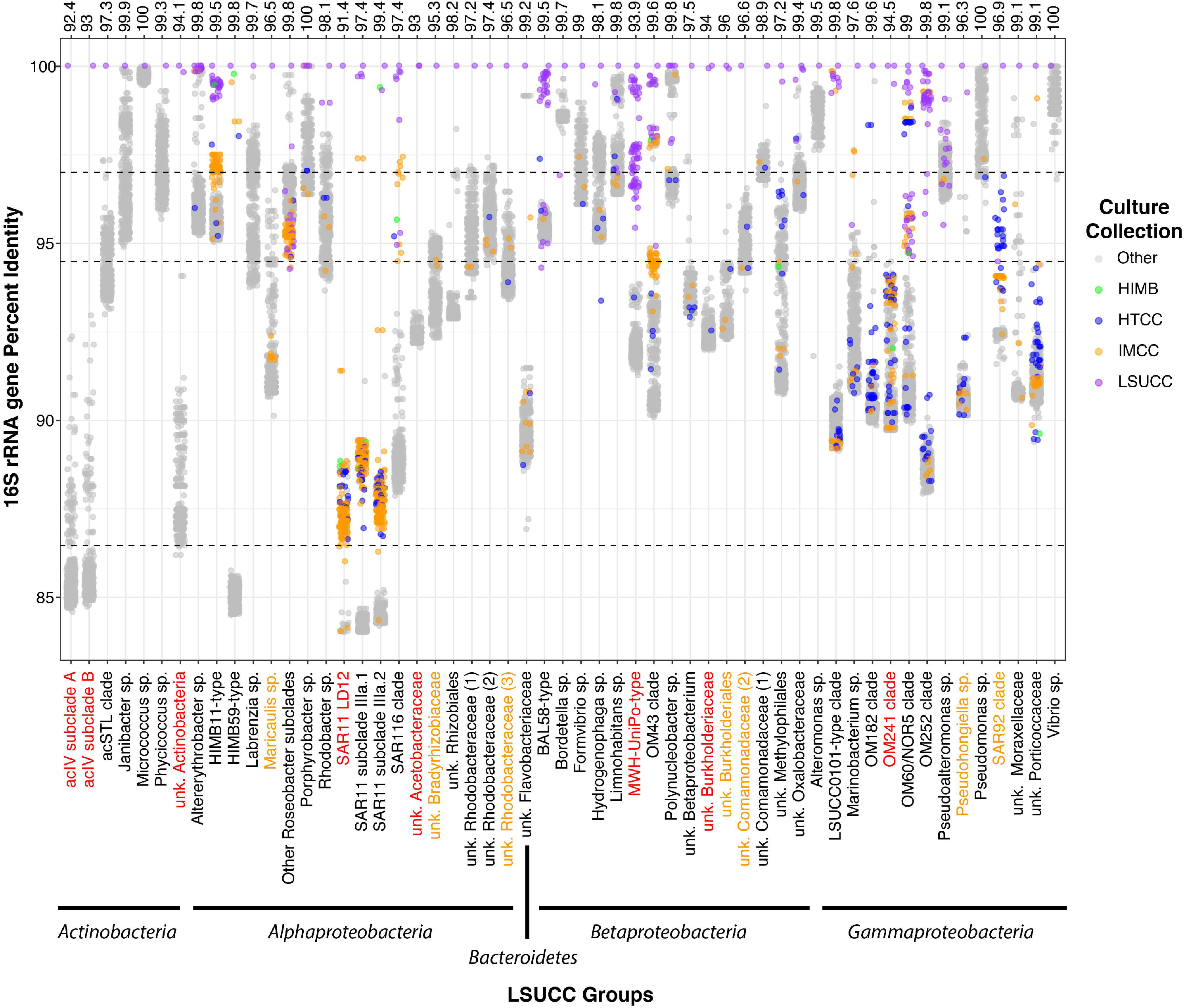
Percent identity of LSUCC isolate 16S rRNA genes compared with those from other isolates in NCBI (“Other”, gray dots) or from the DTE culture collections IMCC (gold dots), HTCC (blue dots), and HIMB (green dots). Each dot represents a pairwise 16S rRNA gene comparison (via BLASTn). X-axis categories are groups designated according to ≥ 94% sequence identity and phylogenetic placement (see Figs S1-S4). Above the graph is the 16S rRNA gene sequence percent identity to the closest non-LSUCC isolate within a column. Groups colored in red indicate those where LSUCC isolates represent putatively novel genera, whereas orange indicates putatively novel species.

### Modeling DTE cultivation via Monte Carlo simulations

We developed a model using Monte Carlo simulation to estimate the median number of positive and pure wells (and associated 95% confidence intervals (CI)) expected from a DTE experiment for a given taxon at different inoculum sizes (*λ*), relative abundances (*r*), and viability (*V*) (Fig. 5). For each bootstrap, the number of cells added to each well was simulated using a Poisson distribution at a mean inoculum size of *λ* cells per well across *n* wells. The number of cells added to each well that belonged to a specific taxon was then estimated using a binomial distribution where the number of trials was set as the number of cells in a well and the probability of a cell belonging to a specific taxon, *r*, was the relative abundance of its representative ASV in the community analysis. Wells that contained at least one cell of a specific taxon were designated ‘positive’. Wells in which all the cells belonged to a specific taxon were designated as ‘pure’. Finally, the influence of taxon-specific viability on recovery of ‘pure’ wells was simulated using a second binomial distribution, where the number of cells within a ‘pure’ well was used as the number of trials and the probability of growth was a viability score ranging from 0 to 1. For each simulation, 9,999 bootstraps were performed. Code for the model and all simulations is available in the ‘viability_test.py’ at our GitHub repository https://github.com/thrash-lab/viability_test.

### Actual versus expected number of isolates

For each taxon in each DTE experiment, the Monte Carlo simulation was used to evaluate whether the number of recovered pure wells for each taxon was within 95% CI of simulated estimates, assuming optimum growth conditions (i.e. *V* = 100%). For each of 9,999 bootstraps, 460 wells were simulated with the inoculum size used for the experiment and the relative abundance of the ASV. For taxa where the number of expected wells fell outside the 95% CI of the model, a deviance score was calculated as the difference between the actual number of wells observed and median of the simulated dataset. The results of this output are presented in Table S1 under the “Expected vs actual” tab, and the R script for visualizing this output as Figure 7 is available at our GitHub repository https://github.com/thrash-lab/EvsA-visualization.

### Estimating viability in under-represented taxa

For taxa where the observed number of positive wells was lower than the 95% CI lower limit within a given experiment, and because our analysis was restricted to only those organisms for which our media was sufficient for growth at least once, the deviance was assumed to be a function of a viability term, *V,* (ranging from 0 to 1) associated with suboptimal growth conditions, dormancy, persister cells, etc. To estimate a value of viability for a given taxon within a particular experiment, the Monte Carlo simulation was run using an experiment-appropriate inoculum size, relative abundance, and number of wells (460 for each experiment). Taxon-specific viability was tested across a range of decreasing values from 99% to 1% until such time as the observed number of pure wells for a given taxon fell between the 95% CI bounds of the simulated data. At this point, the viability value is the maximum viability of the taxon that enables the observed number of pure wells for a given taxon to be explained by the model. The results of this output are presented in Table S1 under the “Expected vs actual” tab.

### Likelihood of recovering taxa at different relative abundances

To estimate the number of wells required in a DTE experiment to have a significant chance of recovering a taxon with a relative abundance of *r,* assuming optimum growth conditions (*V* = 100%), the Monte Carlo model was used to simulate experiments from 92 wells to 9,200 wells per experiment across a range of relative abundances from 0 to 100% in 0.1% increments, and a range of inoculum sizes (cells per well of 1, 1.27, 1.96, 2, 3, 4 and 5). Each experiment was bootstrapped 999 times and the number of bootstraps in which the lower-bound of the 95% CI was ≥ 1 was recorded.

### Data accessibility

All iTag sequences are available at the Short Read Archive with accession numbers SRR6235382-SRR6235415 (29). PCR-generated 16S rRNA gene sequences from this study are accessible on NCBI GenBank under the accession numbers MK603525-MK603769. Previously generated 16S rRNA genes sequences are accessible on NCBI GenBank under the accession numbers KU382357-KU38243 (35). Table S1 is available at https://doi.org/10.6084/m9.figshare.12142113.

## Results

### General cultivation campaign results

We conducted a total of seventeen DTE cultivation experiments to isolate bacterioplankton (sub 2.7 μm fraction), with paired microbial community characterization of source waters (0.22 μm - 2.7 μm fraction), from six coastal Louisiana sites over a three-year period (Table S1). We inoculated 7,820 distinct cultivation wells (all experiments) with an estimated 1-3 cells·well^−1^ using overlapping suites of artificial seawater media, JW (years 1 and 2, (35)) and MWH (year 3), designed to match the natural environment (Table 1). The MWH suite of media was modified from the JW media to additionally include choline, glycerol, glycine betaine, cyanate, DMSO, DMSP, thiosulfate, and orthophosphate (Table S1). These compounds have been identified as important metabolites and osmolytes for marine and freshwater microorganisms and were absent in the first iteration (JW) of our media (88–94). A total of 1,463 wells were positive (> 10^4^ cells·ml^−1^), and 738 of these were transferred to 125 mL polycarbonate flasks. For four experiments (FWC, FWC2, JLB2, and JLB3) we only transferred a subset of positives (48/301, 60/403, 60/103, and 60/146) because the number of isolates exceeded our ability to maintain and identify them at that time (Table 1). The subset of positive wells for these four experiments was selected using flow cytometry signatures usually indicative of smaller oligotrophic cells like SAR11 strain HTCC1062 (49) using our settings. Of the 738 wells from which we transferred cells across all experiments, 328 of these yielded repeatably transferable isolates that we deemed as pure cultures based on 16S rRNA gene PCR and Sanger sequencing.

### Phylogenetic and geographic novelty of our isolates

The 328 isolates belonged to three Phyla: *Proteobacteria* (n = 319), *Actinobacteria* (n = 8), and *Bacteroidetes* (n = 1) (Figs. S1-S5). We placed these isolates into 55 groups based on their positions within 16S rRNA gene phylogenetic trees (Figs. S1-S5) and as a result of having ≥ 94% 16S rRNA gene sequence identity to other isolates. We applied a nomenclature to each group based on previous 16S rRNA gene database designations and/or other cultured representatives (Fig. 1, Table S1). Isolates represented eight putatively novel genera with ≤ 94.5% 16S rRNA gene identity to a previously cultured representative: the *Actinobacteria* acIV subclades A and B, and one other unnamed *Actinobacteria* group; an undescribed *Acetobacteraceae* clade (*Alphaproteobacteria*); the freshwater SAR11 LD12 (*Candidatus* Fonsibacter ubiquis (29)); the MWH-UniPo and an unnamed *Burkholderaceae* clade (*Betaproteobacteria*); and the OM241 *Gammaproteobacteria* (Fig. 1, Table S1). Seven additional putatively novel species in other genera were also isolated (between 94.6 and 96.9% 16S rRNA gene sequence identity) in unnamed *Commamonadaceae* and *Burkholderiales* clades (*Betaproteobacteria*); the SAR92 clade and *Pseudohonigella* genus (*Gammaproteobacteria*); and unnamed *Rhodobacteraceae* and *Bradyrhizobiaceae* clades, as well as *Maricaulis* spp. (*Alphaproteobacteria*) (Fig. 1). LSUCC isolates belonging to the groups BAL58 *Betaproteobacteria* (Fig. S4), OM252 *Gammaproteobacteria*, HIMB59 *Alphaproteobacteria*, and what we designated the LSUCC0101-type *Gammaproteobacteria* (Fig. S5) had close 16S rRNA gene matches to other isolates at the species level, however, none of those previously cultivated organisms have been formally described (Fig. 1). The OM252, BAL58, and MWH-UniPo clades were the most frequently cultivated, with 124 of our 328 isolates belonging to these three groups (Table S1). In total, 73 and 10 of the 328 isolates belonged in putatively novel genera and novel species in previously cultivated genera, respectively. We estimated that at least 310 of these isolates were geographically novel, being the first of their type cultivated from the nGOM (Fig. 2). This included isolates from cosmopolitan groups like SAR11 subclade IIIa, OM43 *Betaproteobacteria*, SAR116, and HIMB11-type “Roseobacter” spp. Cultivars from *Vibrio sp.* and *Alteromonas sp.* were the only two groups with close relatives (species level) isolated from the GOM.

**Figure 2.**
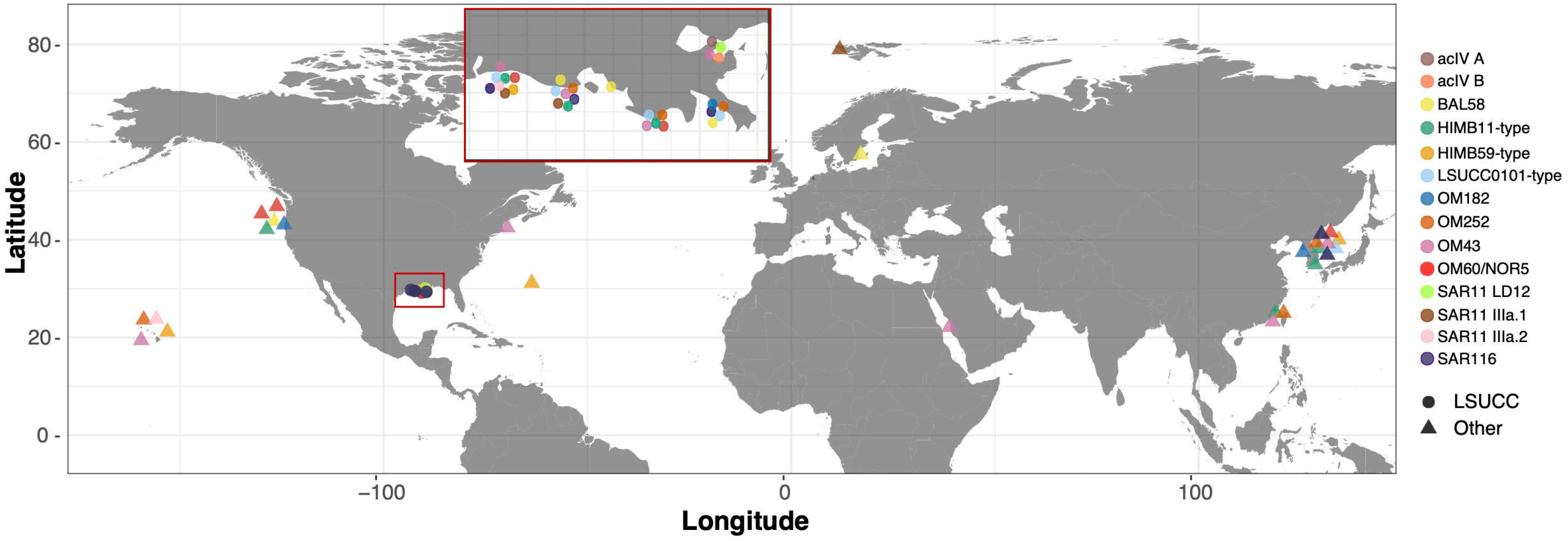
A global map of the isolation location of isolates from selected important aquatic bacterioplankton clades. All depicted taxa were isolated from surface water (< 1-20m), or the depth of sample was not reported (see Table S1 for details). Circles represent LSUCC isolates, while triangles are non-LSUCC isolates. Inset: a zoomed view of the coastal Louisiana region where LSUCC bacterioplankton originated.

### Natural abundance of isolates

We matched LSUCC isolate 16S rRNA gene sequences with both operational taxonomic units (OTUs) and amplicon single nucleotide variants (ASVs) from bacterioplankton communities to assess the relative abundances of our isolates in the coastal nGOM waters that served as inocula. While OTUs provide a broad group-level designation (97% sequence identity), this approach can artificially combine multiple ecologically distinct taxa (95). Due to higher stringency for defining a matching 16S rRNA gene, ASVs can increase the confidence that our isolates represent environmentally relevant organisms (74, 96). However, while many abundant oligotrophic bacterioplankton clades such as SAR11 (29, 97), OM43 (40, 41), SAR116 (98), and *Sphingomonas* spp. (99) have a single copy of the rRNA gene operon, other taxa can have multiple rRNA gene copies (97, 100), complicating ASV analyses. Since we could not *a priori* rule out multiple rRNA gene operons for novel groups with no genome sequenced representatives, we used both OTU and ASV approaches.

In total, we obtained at least one isolate from 40 of the 777 OTUs and 71 of the 1,323 ASVs observed throughout the three-year dataset. 43% and 26% of LSUCC isolates matched the top 50 most abundant OTUs (median relative abundances, all sites, from 8.1-0.11%; Fig. S6A) and ASVs (mean relative abundances, all sites, from 3.8-0.11%; Fig. S6B), respectively, across all sites and samples. Microbial communities from all collected samples clustered into two groups corresponding to those inhabiting salinities below 7 and above 12, and salinity was the primary environmental driver distinguishing community beta diversity (OTU: R^2^=0.88, P=0.001, ASV: R^2^=0.89, P =0.001). As part of the cultivation strategy after the first five experiments, we used a suite of five media differing by salinity and matched the experiment with the medium that most closely resembled the salinity at the sample site. Consequently, our isolates matched abundant environmental groups from both high and low salinity regimes. At salinities above twelve, LSUCC isolates matched 13 and 14 of the 50 most abundant OTUs and ASVs, respectively (Figs. 3A, 4A; Table S1). These taxa included the abundant SAR11 subclade IIIa.1, HIMB59, HIMB11-type “Roseobacter”, and SAR116 *Alphaproteobacteria*; the OM43 *Betaproteobacteria*; and the OM182 and LSUCC0101-type *Gammaproteobacteria*. At salinities below seven, 10 and 9 of the 50 most abundant OTUs and ASVs, respectively, were represented by LSUCC isolates, including one of most abundant taxa in both cluster sets, SAR11 LD12 (Figs. 3B, 4B). Some taxa, such as SAR11 IIIa.1 and OM43, were among the top 15 most abundant taxa in both salinity regimes (Figs. 3, 4, Table S1), suggesting a euryhaline lifestyle. In fact, our cultured SAR11 IIIa.1 ASV7471 was the most abundant ASV in the aggregate dataset (Fig. S6).

**Figure 3.**
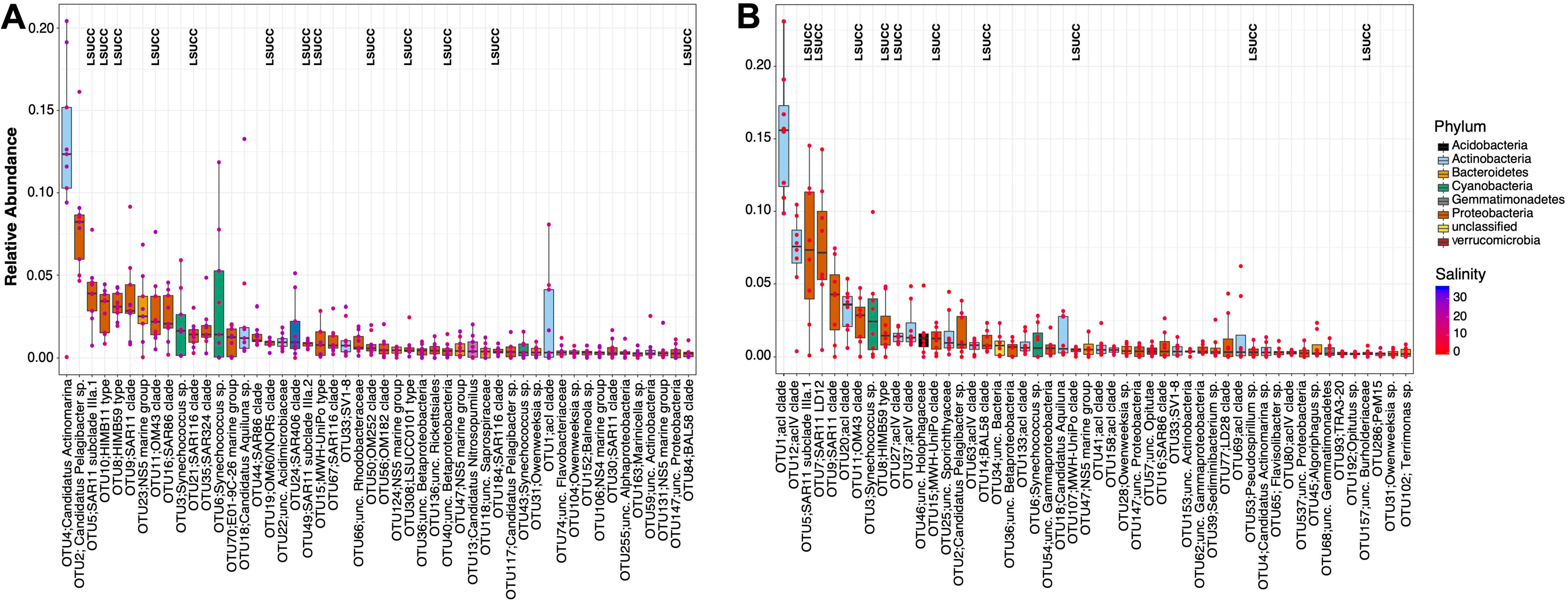
Rank abundances of the 50 most abundant OTUs from all sites based on median relative abundance at salinities less than seven (A) and greater than twelve (B). The boxes indicate the interquartile range (IQR) of the data, with vertical lines indicating the upper and lower extremes according to 1.5 × IQR. Horizontal lines within each box indicate the median. The data points comprising the distribution are plotted on top of the boxplots. The shade of the dot represents the salinity at the sample site (red-blue :: lower-higher), while the color of the box indicates broad taxonomic identity. LSUCC labels indicate OTUs with at least one cultivated representative.

**Figure 4.**
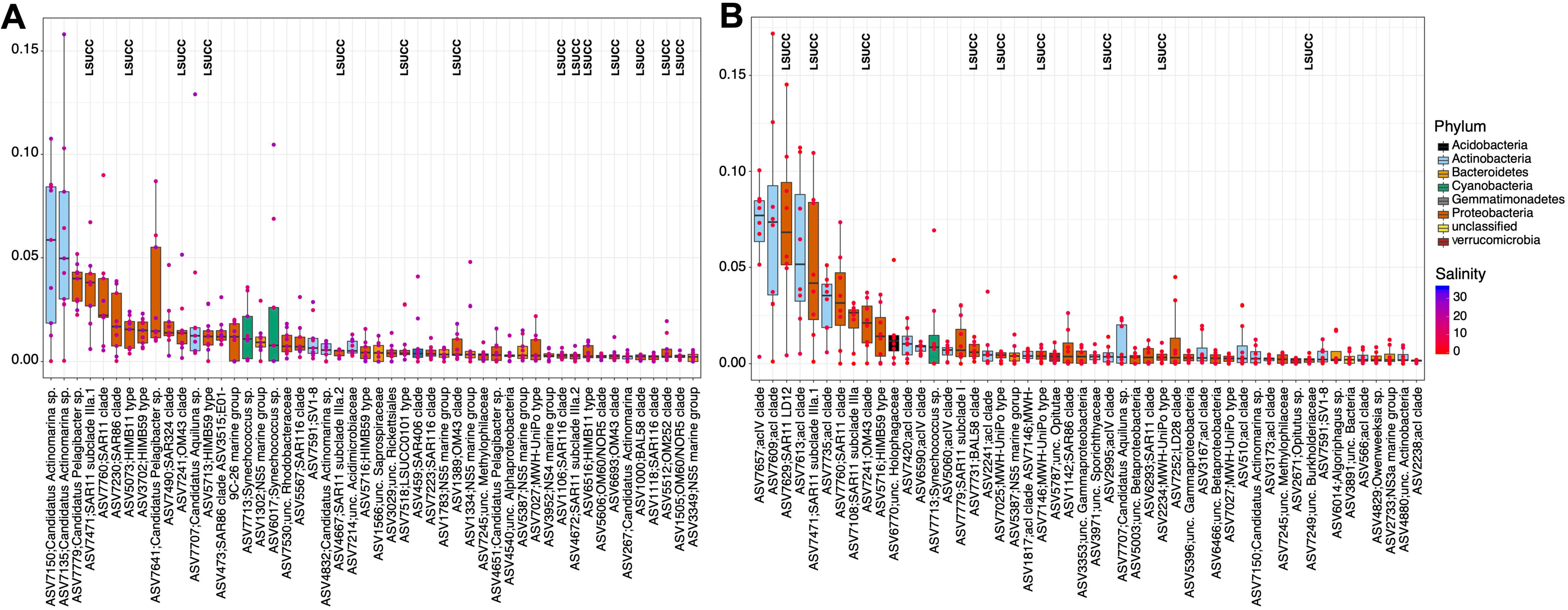
Rank abundances of the 50 most abundant ASVs from all sites based on median relative abundance at salinities less than seven (A) and greater than twelve (B). The boxes indicate the interquartile range (IQR) of the data, with vertical lines indicating the upper and lower extremes according to 1.5 × IQR. Horizontal lines within each box indicate the median. The data points comprising the distribution are plotted on top of the boxplots. The shade of the dot represents the salinity at the sample site (red-blue :: lower-higher), while the color of the box indicates broad taxonomic identity. LSUCC labels indicate ASVs with at least one cultivated representative.

**Figure 5.**
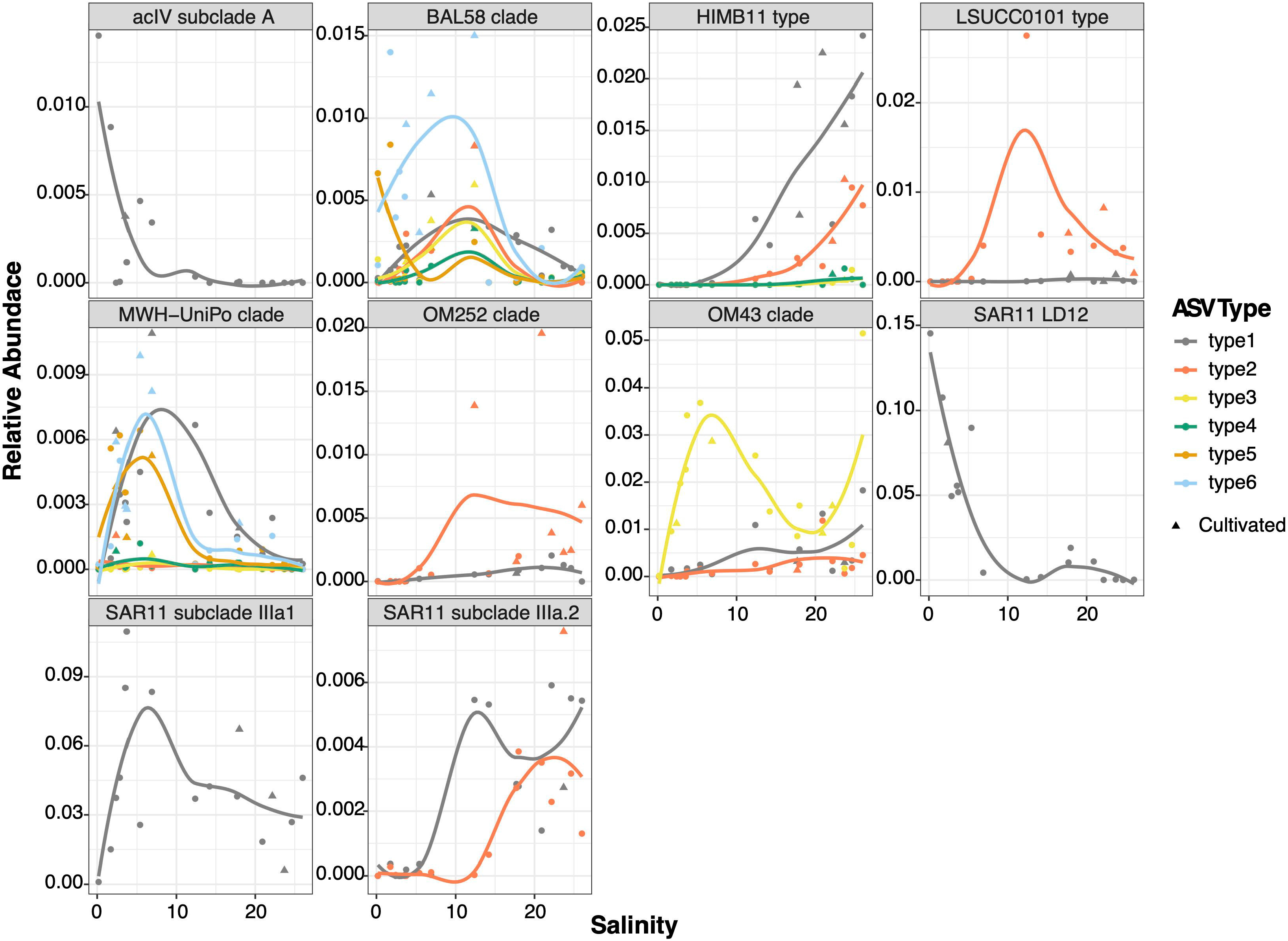
Relative abundance of ASVs within key taxonomic groups compared with salinity. ASV types are colored independently, and triangle points indicate experiments for which at least one isolate was obtained. Non-linear regression lines are provided as a visual aid for abundance trends.

Overall, this effort isolated taxa representing 18 and 12 of the top 50 most abundant OTUs and ASVs, respectively (Table 2, Fig. S6). When looking at different median relative abundance categories of > 1%, 0.1% - 1%, and < 0.1%, isolate OTUs were distributed across those categories in the following percentages: 15%, 20%, and 27%; isolate ASVs were distributed accordingly: 4%, 26%, and 37% (Table 2). Isolates with median relative abundances of < 0.1%, such as *Pseudohongiella* spp., *Rhodobacter* spp., and *Bordetella* spp., would canonically fall within the rare biosphere (101) (Table S1). A number of isolates did not match any identified OTUs or ASVs (38% and 33% of LSUCC isolates when compared to available OTUs and ASVs, respectively), either because their matching OTUs/ASVs were below our thresholds for inclusion (at least two reads from at least two sites), or because they were below the detection limit from our sequencing effort (Table 2). Thus, 43% and 30% of our isolates belonged to OTUs and ASVs, respectively, with median relative abundances > 0.1%.

**Table 2.** Median relative abundances (*r*) of cultured OTUs and ASVs across all samples

| | In top 50 ranks | $r > 1\%$ | $1\% - 0.1\% r$ | $r < 0.1\%$ | Not detected |
| --- | --- | --- | --- | --- | --- |
| OTUs | 18 (140 isolates) | 50 isolates (15%) | 90 isolates (27%) | 65 isolates (20%) | 123 isolates (38%) |
| ASVs | 12 (84 isolates) | 13 isolates (4%) | 84 isolates (26%) | 122 isolates (37%) | 109 isolates (33%) |

### Modeling DTE cultivation

An enigma that became immediately apparent through a review of our data was the absence of an obvious relationship between the abundance of a given taxon in the inoculum and the frequency of obtaining an isolate of the same type from a DTE cultivation experiment (Figs. S7, S8). For example, although we could culture SAR11 LD12 over a range of media conditions (29), and the matching ASV had relative abundances of > 5% in six of our seventeen experiments (Fig. 5), we only isolated one representative (LSUCC0530). In an ideal DTE cultivation experiment where cells are randomly subsampled from a Poisson-distributed population, if the medium is sufficient for a given microorganism’s growth, then the number of isolates should correlate with that microorganism’s abundance in the inoculum. However, a qualitative examination of several abundant taxa that grew in our media, some of which we cultured on multiple occasions, revealed no clear pattern between abundance and isolation success (Fig. 5). Considering that medium composition was sufficient for cultivation of these organisms on at least some occasions, we hypothesized that cultivation frequency may reflect differences in the capacity for growth within populations of a given taxon. Thus, we decided to model cultivation frequency in relationship to estimated abundances in a way that could generate estimates of cellular viability, defined herein as meaning “presently able to grow in defined medium,” as opposed to a broader definition equating viability with being alive more generally, since we only evaluated growth capacity in this study. We hoped that modeling might also help us inform experimental design and make DTE cultivation efforts more predictable (59).

Previously, Don Button and colleagues developed a statistical model for viability (*V*) of cells in the entire population for a DTE experiment (33):

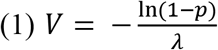

Where *p* is the proportion of wells or tubes, *n*, with growth, *z*, (*p* = *z*/*n*) and *λ* is the estimated number of cells inoculated per well (the authors used *X* originally). The equation uses a Poisson distribution to account for the variability in cell distribution within the inoculum and therefore the variability in the number of wells or tubes receiving the expected number of cells. We and others have used this equation in the past (26, 28, 35) to evaluate the efficacy of our cultivation experiments in the context of commonly cited numbers for cultivability using agar-plate based methods (13, 17, 102).

While Equation 1 was effective for its intended purpose, it has a number of drawbacks that limit its utility for taxon-specific application: 1) If *p=1*, i.e. all wells are positive, then the equation is invalid; 2) At high values of *p* and low values of *λ*, estimates of *V* can exceed 100% (Table 1); 3) Accuracy of viability, calculated by the asymptotic standard error, *ASE*, or the coefficient of variation, *CVV*, was shown to be non-uniform across a range of *λ*, with greatest accuracy when true viability was ~10% (33). Thus, low viability, low values of *λ,* and small values of *n* were found to produce unreliable results; 4) If *p=0*, i.e. no positive wells are observed, estimates of viability that could produce 0 positive wells cannot be calculated. In addition, 5) Button’s original model assumes that a well will only produce a pure culture if the inoculated well contains one cell. By contrast, in low diversity samples, samples dominated by a single taxon, or experiments evaluating viability from axenic cultures across different media, a limitation that only wells with single cells are axenic will underestimate the expected number of pure wells.

To overcome these limitations, we developed a Monte Carlo simulation model that facilitates the incorporation of relative abundance data from complementary community profiling data (e.g. 16S rRNA gene amplicons) to calculate the likelihood of positive wells, pure wells, and viability at a taxon-specific level, based on the observed number of wells for which we obtained an isolate of a particular taxon (Fig. 6). By employing a Monte Carlo approach, our model is robust across all values of *p* and *n* with uniform prediction accuracy, and we can estimate the accuracy of our prediction within 95% confidence intervals (CI). Furthermore, the width of 95% CI boundaries of viability, as well as the expected number of positive and pure wells, are entirely controllable and dependent only on available computational capacity for bootstrapping (i.e., these can be improved with more bootstrapping, but at greater computational cost). When zero positive wells are observed experimentally, our approach enables estimation of a maximum viability that could explain such an observation by identifying the range of variability values for which zero resides within the bootstrapped 95% CI. Finally, the ability to calculate the viability of the entire community, as in Equation 1, is retained simply by estimating viability using a relative abundance of one.

**Figure 6.**
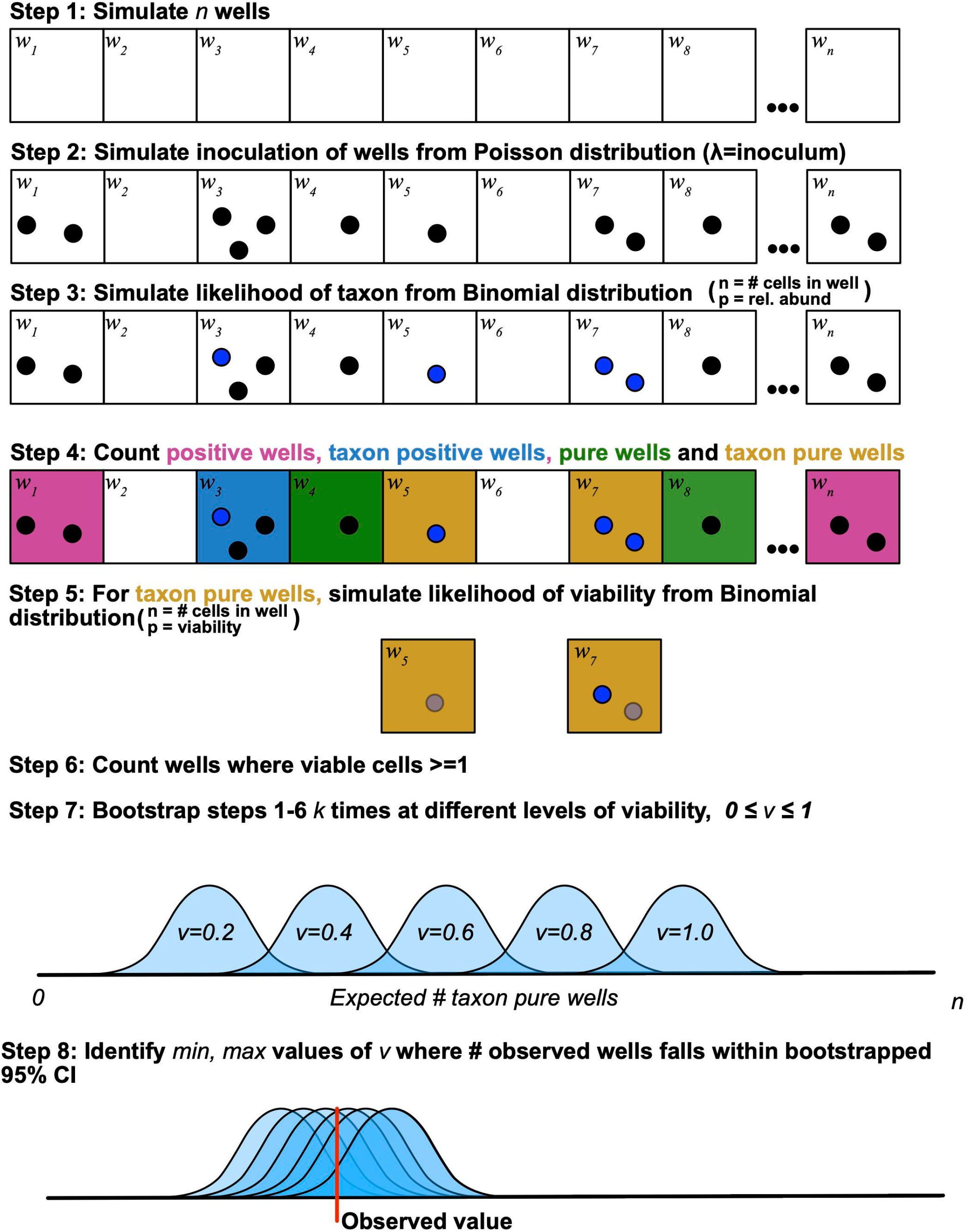
Graphical depiction of the viability model.

**Figure 7.**
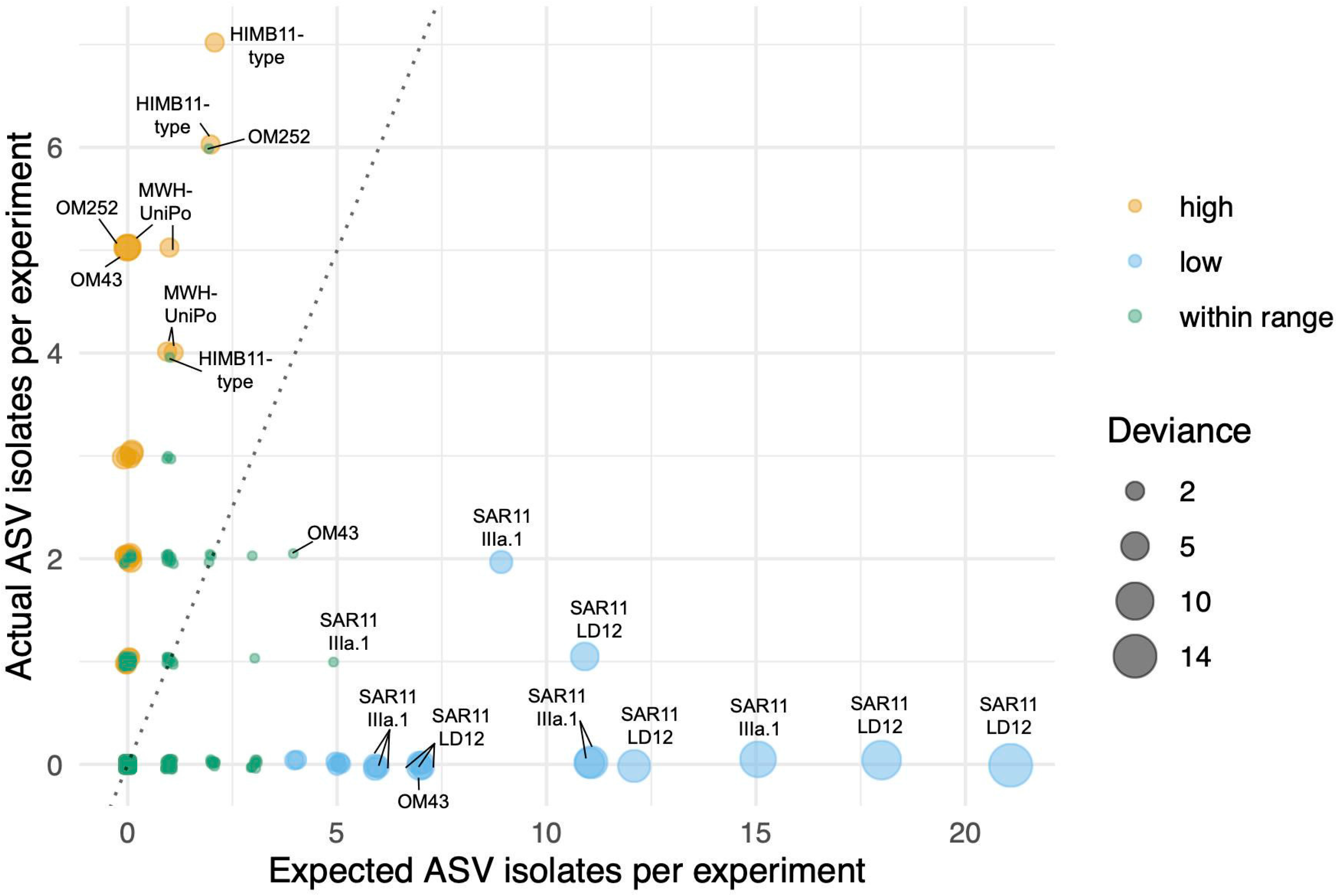
Actual vs. expected numbers of isolates. Each point represents the actual number of isolates for every ASV/experiment pair compared to the expected number of isolates based on our model assuming 100% viability. Colors represent the relationship to the model predictions: green-isolates within the 95% CI for expected frequency, orange-actual isolates > maximum 95% CI for expected isolates; blue-actual isolates < minimum 95% CI for expected isolates. Circle size is proportional to the deviation of the number of actual isolates from the maximum (for orange) or minimum (for blue) 95% CI for expected isolates. The dotted line is the 1:1 ratio. Notable taxa on the extremities of the actual and expected values are labeled. All datapoints provided in Table S1.

We compared our model to that of Button et al. for evaluating viability from whole community experimental results, similarly to previous reports (26, 28, 35) (Table 1). Our viability estimates (*V_est_*) generally agreed with those using the Button et al. calculation, but we have now provided 95% CI to depict the maximum and minimum viability that would match the returned positive well distribution, as well as maximum and minimum values for the number of wells that ought to have contained a single cell. Maximum *V_est_* ranged from 1.1% to > 92.3% depending on the experiment, with a median *V_est_* across all our experiments of 8.6% (Table 1). In one case, the extremely high value (FWC2) was better handled by our model compared to equation 1, because it did not lead to a viability estimation greater than 100%. FWC and FWC2 represent *V_est_* outliers compared with the entire dataset (maximums of 59.7% and > 92.3%, respectively; Table 1). We believe these high numbers most likely resulted from underestimating the number of cells inoculated into each well (because of the use of microscopy, the presence of clumped cells, or possible pipet error-described in (35)), thus increasing the estimated viability.

### Isolate-specific viability estimates

Our new model also facilitates taxon-specific viability estimates. Cultivation efficacy was evaluated for 71 cultured taxa matching ASVs within our detection limits (219 isolates) across 17 sites (1,207 pairwise combinations) by comparing the number of observed pure wells to those predicted by the Monte Carlo simulation using 9,999 bootstraps, 460 wells per experiment, and an assumption that all cells were viable (i.e. *V* = 100%). In total, for 1,158 out of 1,207 pairwise combinations (95.9%) the observed number of pure wells fell within the 95% CI of data simulated at matching relative abundance and inoculum size, suggesting that these two parameters alone could explain the observed cultivation success for most taxa (Table S1). 1,059 out of these 1,158 combinations (91%) recorded zero observed wells, but with a maximum relative abundance of 2.8% within these combinations, a score of zero fell within predicted 95% CI of simulations with 460 wells. Sensitivity analysis showed that with 460 wells per experiment, an observation of zero pure wells falls below the 95% confidence intervals lower-bound (and is thus significantly depleted to enable viability to be estimated) for taxa with relative abundances of 2.3%, 2.9% and 4.5% for inoculum sizes of one, two and three cells per well, respectively (Fig. S9). In fact, modeling DTE experiments from 92 wells to 9,200 wells per experiment showed that for taxa comprising ~1% of a microbial community 1,104 wells (or 12 plates at 92 wells per plate), 1,380 wells (15 plates) and 2,576 wells (28 plates) were required to be statistically likely to recover at least one positive, pure well using inocula of one, two or three cells per well, respectively, with *V* = 100% (Fig. S9).

A small, but taxonomically relevant minority (49 out of 1,207) of pairwise combinations had a number of observed pure wells that fell outside of the simulated 95% CI with *V* = 100% (Fig. 7). Of these, 28 had either one, two, or three more observed pure wells than the upper 95% CI (Table S1), suggesting cultivability *higher* than expected based purely a model capturing the interaction between a Poisson-distributed inoculum and a binomially-distributed relative abundance, with *V* = 100%. However, the deviance from the expected number of positive wells for those above the 95% CI was limited to three or fewer wells, meaning that we only obtained 1-3 more isolates than expected (Table S1). On the other hand, those organisms that we isolated less frequently than expected showed greater deviance. 21 out of the 49 outliers had lower than expected cultivability (Fig. 7). These taxa had relative abundances ranging from 2.7% to 14.5%, but recorded only 0, 1, or 2 isolates. In the most extreme case, ASV7629 (SAR11 LD12) at Site ARD2c comprised 14.5% of the community but recorded no observed pure wells, compared to expected number of 13-30 isolates (95% CI) predicted by the Monte Carlo simulation. All the examples of taxa that were isolated less frequently than expected given the assumption of *V* = 100% belonged to either SAR11 LD12, SAR11 IIIa.1, or one particular OM43 ASV (7241) (Fig. 6).

We used our model to calculate estimated viability (*V_est_*) for these organisms based on their cultivation frequency at sites where the assumption of *V* = 100% appeared violated (Table 3). Using the extreme example of SAR11 LD12 ASV7629 at Site ARD2c, simulations across a range of *V* indicated that a result of zero positive wells fell within 95% of simulated values when the associated taxon *V_est_* ≤ 15%. When considering all anomalous cultivation results, LD12 had estimated maximum viabilities that ranged up to 55% (Table 3). OM43 (ASV7241) estimated maximum viabilities ranged from 52-80%, depending on the site, and similarly, SAR11 IIIa.1 ranged between 22-82% maximum viability (Table 3).

**Table 3.** Estimated viabilities for taxa cultivated less frequently than expected

| ASV | Group | Site | $r^*$ | $n$ | $z$ | $\lambda$ | Estimated # wells with 1 cell (bootstrapped median: (xx-xx) 95% CI) if $V=1$ ** | $V_{est}$: min-max 95% CI based on cultivation results*** |
| --- | --- | --- | --- | --- | --- | --- | --- | --- |
| 7241 | OM43 | ARD3 | 0.03 | 460 | 0 | 2 | 4 (1-9) | 0.1-80 |
| 7241 | OM43 | FWC<sup>†</sup> | 0.04 | 460 | 0 | 2 | 5 (1-9) | 0.1-77 |
| 7241 | OM43 | JLB | 0.05 | 460 | 0 | 1.96 | 7 (2-12) | 0.1-52 |
| 7471 | SAR11 IIIa.1 | ARD3 | 0.11 | 460 | 0 | 2 | 15 (8-23) | 0.1-22 |
| 7471 | SAR11 IIIa.1 | CJ | 0.03 | 460 | 0 | 1.27 | 4 (1-9) | 0.1-82 |
| 7471 | SAR11 IIIa.1 | FWC3 | 0.07 | 460 | 2 | 2 | 9 (4-15) | 2.5-80 |
| 7471 | SAR11 IIIa.1 | JLB | 0.05 | 460 | 0 | 1.96 | 6 (2-11) | 0.1-59 |
| 7471 | SAR11 IIIa.1 | JLB2c<sup>†</sup> | 0.08 | 460 | 0 | 2 | 11 (5-18) | 0.1-31 |
| 7471 | SAR11 IIIa.1 | JLB3<sup>†</sup> | 0.04 | 460 | 0 | 2 | 5 (1-9) | 0.1-74 |
| 7471 | SAR11 IIIa.1 | LKB | 0.05 | 460 | 0 | 1.8 | 6 (2-12) | 0.1-55 |
| 7471 | SAR11 IIIa.1 | LKB2 | 0.04 | 460 | 0 | 2 | 5 (1-9) | 0.1-77 |
| 7471 | SAR11 IIIa.1 | LKB3 | 0.09 | 460 | 0 | 2 | 11 (6-19) | 0.1-30 |
| 7471 | SAR11 IIIa.1 | TBON2 | 0.04 | 460 | 0 | 1.56 | 6 (2-12) | 0.1-56 |
| 7471 | SAR11 IIIa.1 | TBON3 | 0.04 | 460 | 0 | 2 | 5 (1-10) | 0.1-73 |
| 7629 | SAR11 LD12 | ARD | 0.11 | 460 | 0 | 1.5 | 18 (10-27) | 0.1-20 |
| 7629 | SAR11 LD12 | ARD2c | 0.15 | 460 | 0 | 2 | 21 (13-30) | 0.1-15 |
| 7629 | SAR11 LD12 | ARD3 | 0.05 | 460 | 0 | 2 | 7 (2-12) | 0.1-53 |
| 7629 | SAR11 LD12 | FWC<sup>†</sup> | 0.09 | 460 | 0 | 2 | 12 (6-19) | 0.1-28 |
| 7629 | SAR11 LD12 | LKB | 0.05 | 460 | 0 | 1.8 | 7 (2-13) | 0.1-51 |
| 7629 | SAR11 LD12 | LKB2 | 0.08 | 460 | 1 | 2 | 11 (5-18) | 0.3-49 |
| 7629 | SAR11 LD12 | LKB3 | 0.06 | 460 | 0 | 2 | 7 (3-13) | 0.1-48 |

\*Fractional relative abundance
\*\*Based on 9,999 bootstraps.
\*\*\*Based on 9,999 bootstraps tested at viability increments of 0.1%.
<sup>†</sup>Experiments where a subset of positive wells were transferred.

## Discussion

This work paired 17 DTE cultivation experiments with cultivation-independent assessments of microbial community structure in source waters to evaluate cultivation efficacy. We generated 328 new bacterial isolates representing 40 of the 777 OTUs and 71 of the 1,323 ASVs observed across all samples from which we inoculated DTE experiments. Stated another way, we successfully cultivated 5% of the total three year bacterioplankton community observed via either OTU or ASV analyses. A large fraction of our isolates (43% of cultured OTUs, 30% of cultured ASVs) represented taxa present at median relative abundances > 0.1%, with 15% and 4% of cultured OTUs and ASVs, respectively, at median abundances > 1%. 140 of our isolates matched the top 50 most abundant OTUs, and 84 isolates matched the top 50 most abundant ASVs.

This campaign led to the first isolations of the abundant SAR11 LD12 and *Actinobacteria* acIV; the second isolate of the HIMB59 *Alphaproteobacteria*; and new genera within the *Acetobacteraceae*, *Burkholderiaceae*, OM241 and LSUCC0101-type *Gammaproteobacteria*, and MWH-UniPo *Betaproteobacteria*; thereby demonstrating again that continued DTE experimentation leads to isolation of previously uncultured organisms with value for aquatic microbiology. We have also added a considerable collection of isolates to previously cultured groups like OM252 *Gammaproteobacteria*, BAL58 *Betaproteobacteria*, and HIMB11-type “Roseobacter” spp., and the majority of our isolates represent the first versions of these types of taxa from the Gulf of Mexico, adding comparative biogeographic value to these cultures.

Our viability model improved upon the statistical equation developed by Button and colleagues (33) to extend viability estimates to individual taxa within a mixed community and provide 95% CI constraining those viability estimates. We cultured several groups of organisms abundant enough to evaluate viability with 460 wells (Figs. 7, S9). The fact that these organisms were successfully cultured at least once meant that we could reasonably assume that the medium was sufficient for growth. Some taxa were cultivated more frequently than expected (Fig. 7). We explore two possible explanations for this phenomenon-errors in quantification and variation in microbial cell organization. Any systematic error that led to underestimating the abundance of an organism would have correspondingly resulted in our underestimating the number of wells in which we would expect to find a pure culture of that organism. Such underestimations could come from primer biases associated with amplicon sequencing (69, 70, 103), but we do not know if those protocols specifically underestimate the OM252, MWH-UniPo, and HIMB11-type taxa cultured more frequently than expected (Fig. 7). However, due the low number of expected isolates in these groups, and the small deviances in actual isolates from those expected numbers (within 1-3 isolates compared to expected values), the biases inherent in the relative abundance estimations for these taxa were probably small. Furthermore, one of the microorganisms isolated more frequently than expected matched the OM43 ASV1389 (Fig. S6), whereas another OM43 ASV (7241) was cultivated less frequently than expected (see below), meaning that if primer bias were the cause of this discrepancy, it would have to be operating differently on very closely related organisms.

One possible biological explanation for why some isolates might have been cultured more frequently than expected is clumped cells. If cells of any given taxon in nature grew in small clusters, then the number of cells we added to a well would have been greater than expected based on a Poisson distribution. Furthermore, the model assumes that each cell is independent, and that the composition of a subset of cells is only a function of the relative abundance of the taxon in the community. Within a cluster of cells, this assumption is violated as the probability of cells being from the same taxon is higher. Thus, the model will underestimate the probability of a well being pure and therefore underestimate the number of pure wells likely to be observed within an experiment, leading to a greater number of isolates than expected. Future microscopy work could examine whether microorganisms such as OM252 and MWH-UniPo form small clusters *in situ* and/or in pure culture, and whether this phenomenon may be different for different ASVs of OM43, or if clumping may be a transient phenotype.

We also identified three taxa-SAR11 LD12, SAR11 subclade IIIa.1, and the aforementioned OM43 ASV7241-that were isolated much less frequently than expected based on their abundances (Fig. 7, Table 3). This could mean that our assumption of *V* = 100% was incorrect, or that, in contrast to the taxa that were cultured more frequently than expected (above), our methods had biases that *over*estimated the abundance of these organisms, thereby over-inflating the expected number of isolates. We used the modified 515/806RB primers that have been shown to be much more accurate in quantifying SAR11 compared to FISH than the original 515/806 primers (within 6% ± 4% SD), and this protocol almost always underestimates SAR11 abundance (69). This suggests that our expected number of isolates may have actually been underestimated, our cultivation success poorer than we measured, and therefore we may be overestimating viability for the SAR11 taxa in this study. Other sources of systematic error that might impinge on successful transfers, and thereby reduce our recovery, include sensitivity to pipette tip and/or flask material. However, the fact that these taxa were sometimes successfully isolated means that if these mechanisms were impacting successful transfers, then their activity was less than 100% efficient, which implies variations in subpopulation vulnerability that would be very similar conceptually to variations in subpopulation viability.

Another possible source of error that could have resulted in lower than expected numbers of isolates was the subset of experiments for which we did not transfer all positive wells due to limitations in available personnel time (Tables 1 & 3). However, our selection criteria for the subset of wells to transfer was based on flow cytometric signatures that would have encompassed small cells like SAR11 (see Results), and in any case, there were many examples of lower than expected recovery from other experiments where we transferred all positive wells (Table 3). Thus, we believe that these four experiments were unlikely to contribute major errors biasing our estimates of viability for SAR11 LD12, SAR11 IIIa.1, and other small cells like OM43.

If we instead explore biological reasons for the lower than expected numbers of positive wells in DTE experiments, a plausible explanation supported by the literature is simply that a large fraction of the population is in some state of inactivity or at least not actively dividing (104). Studies using uptake of a variety of radiolabelled carbon and sulfur sources have demonstrated substantial fractions of SAR11 cells may be inactive depending on the population (105–108). SAR11 cells in the northwest Atlantic and Mediterranean showed variable uptake of labelled leucine (30-50% (105, 106); ~25-55% (108, 109)) and amino acids (34-61% (105, 106); 34-66% (105, 106)). Taken in reverse, this means that up to 75% of the SAR11 population may be dormant at any given time. In another study focused on brackish communities, less than 10% of SAR11 LD12 cells took up labelled leucine and/or thymidine (107). While this was likely not the ideal habitat for LD12 based on salinities above six (29, 107), this study supports the others above that show substantial proportions of inactive SAR11 cells, the fraction of which may depend on environmental conditions and other unknown factors. Bi-orthagonal non-canonical amino acid tagging (BONCAT) shows similar trends for SAR11 (110). These results also match general data indicating prevalent inactivity among aquatic bacterioplankton (104, 111–113). Although labelled uptake methods do not directly measure rates of cell division, the incorporation of these compounds requires active DNA replication or translation, which represent an even more fundamental level of activity than cell division (114).

Why might selection favor high percentages of subpopulation dormancy? One possibility is as an effective defense mechanism against abundant viruses. Viruses infecting SAR11 have been shown to be extremely abundant in both marine (115, 116) and freshwater (117) systems. Indeed, the paradox of high viral abundances and high host abundances in SAR11 has led to a refining of negative density dependent selection through Lokta-Volterra predator-prey dynamics (118) to include heterogeneous susceptibility at the strain level (119, 120) and positive density dependent selection through intraspecific proliferation of defense mechanisms (121). Activity of lytic viruses infecting SAR11 *in situ* demonstrated that phages infecting SAR11 have lower ratios of viral transcripts to host cells compared to other abundant taxa, and that observed abrupt changes in these ratios suggest co-existence of several SAR11 strains with different life strategies and phage susceptibility (122). Phenotypic stochasticity of phage receptor expression has been shown to maintain a small proportion of phage-insensitive hosts within a population, enabling coexistence of predator and prey without extinction (123). Phages adsorb to a vast array of receptor proteins on their hosts, with many well-characterised receptors (e.g. OmpC, TonB, BtuB, LamB) associated with nutrient uptake or osmoregulation (124). Selection therefore favours phenotypes that limit receptor expression, with an associated fitness cost, particularly in nutrient-limited environments.

However, an alternative mechanism is possible if a population of cells comprised a small number of susceptible cells, and a large number of either resistant or dormant cells where presentation of receptor proteins is retained. The majority of host-virus encounters would occur with resistant or dormant cells, and constrain viral propagation through inefficient or failed infection, effectively acting as a sink for infectious particles. Prevalent lysogeny in SAR11 populations would provide a mechanism for establishing resistant cells via superinfection immunity (125, 126), where integration of a temperate phage prevents infection by other closely related viruses. There is growing evidence that many viruses infecting SAR11 are temperate (127, 128) and that reversion to virulence can be triggered through nutrient limitation (128) in contrast to other systems where lysogeny is favoured in nutrient-poor conditions (129). Viral infection may also trigger host dormancy, lowering cellular metabolism to minimise energy requirements under nutrient limited conditions (130). Such cells would be selected against during cultivation experiments, potentially explaining the rarity of SAR11 isolate genomes found to contain prophages. Dormancy and/or lysogeny would also enable long-term co-stability between abundant phages and their hosts (131) and resolve the apparent paradox of high host and virus abundances (126).

Detailed measurements of dormancy in SAR11 and what types of cellular functions become inactivated are part of our ongoing work. In the meantime, it is prudent to examine the implications of a substantial proportion of non-dividing cells for our understanding of basic growth dynamics. Studies attempting to measure SAR11 growth rates in nature have yielded a wide range of results, ranging from 0.03-1.8 day^−1^ (97, 105, 108, 132–134). These span wider growth rates than observed for axenic cultures of SAR11 (0.4-1.2 day^−1^), but isolate-specific growth ranges within that spread are much more constrained (29, 36, 49, 135, 136). Conversion factors for determining production from ^3^H-leucine incorporation (137) are accurate for at least some Ia subclade members of SAR11 (138), so variations in growth rate estimates from microradiography experiments likely have other explanations. It is possible that different strains of SAR11 simply have variations in growth rate not captured by existing isolates. Another, not mutually exclusive, possibility is that the differences in *in situ* growth rate estimates also reflect variations in the proportion of actively dividing cells within the population. A simple model of cell division with binary fission where only a subset of cells divide and non-dividing cells persist, rather than die, can still yield logarithmic growth curves (Fig. S10) like those observed for SAR11 in pure culture (29, 49, 139). However, this subpopulation variability means that the division rate for the subset of cells that are actively dividing is much higher than calculated when assuming 100% dividing cells in the population. Based on our estimated viability for SAR11 LD12 of 15-55%, to obtain our previously calculated maximum division rate (0.5 day^−1^) for the whole culture (29), the per-cell division rate for only a subpopulation would span 2.48-0.79 day^−1^ (Fig. S10, Supplemental Text). Verifying the proportion of SAR11 cells actively dividing in a culture may be challenging. Time-lapse microscopy (140) offers an elegant solution if SAR11 can be maintained for the requisite time periods for accurate measurements in a microfluidic device.

In addition to identifying taxa whose isolation success suggested deviations from biological assumptions of single planktonic cells with 100% viability, the model also revealed the limitations of DTE cultivation in assessing viability depending on relative abundance (Fig. S9). We cannot ascertain whether any given taxon may violate an assumption of *V* = 100% unless we have enough wells to demonstrate that it grew in fewer wells than expected. For example, taxa at 1% of the microbial community require more than 1,000 wells before the lack of a cultured organism represents a significant negative event, rather than a taxon simply lacking sufficient abundance to ensure inclusion in a well within 95% CI. In our 460 well experiments, we could not resolve whether taxa may have had viabilities below 100% if they were less than 3% of the community for any given experiment (Fig. S9). Modeling DTE experiments showed that for experiments targeting rare taxa, lower inoculum sizes are favoured where selective media for enrichment is either unknown or undesirable. The exponential increase in the number of required wells with respect to the inoculum size is a function of a pure well requiring *all* cells within it to belong to the same taxon, assuming all cells are equally and optimally viable.

By providing taxon-specific predictions of viability from cultivation data, our model now facilitates an iterative process to improve experimental design and make cultivation more reliable. First, we use the cultivation success rates to determine for which taxa the assumption of 100% viability was violated. Second, we use the model to estimate viability for those organisms. Third, we use the viability and relative abundance data to determine, within 95% CI, the appropriate number of inoculation attempts required to isolate a new version of that organism. Using SAR11 LD12 as an example, given a relative abundance of 10%, and a viability of 15%, 800 DTE wells should yield four pure, positive wells (1-8 95% CI). This means that, for microorganisms that we know successfully grow in our media, we can now statistically constrain the appropriate number of wells required to culture a given taxon again. For organisms that were not abundant enough to estimate viability using the model, we can use a conservative viability assumption (e.g., 50% (111)) with which to base our cultivation strategy, thereby still reducing uncertainty about the experimental effort necessary to re-isolate one of these microorganisms.

## Conclusions

This work has provided hundreds of new cultures for microbiological research, many among the most abundant members of the nGOM coastal bacterioplankton community. It also provides another demonstration of the effectiveness of sustained cultivation efforts for bringing previously uncultivated strains into culture. Our modeled cultivation results have generated compelling evidence for low viability within subpopulations of SAR11 LD12 and IIIa.1, as well as OM43 *Betaproteobacteria*. The prevalence of, and controls on, dormancy in these clades deserves further study. We anticipate that future work with larger DTE experiments will yield similar viability data about other groups of taxa with lower abundance, highlighting a valuable diagnostic application of DTE cultivation/modeling beyond the primary role in isolating new microorganisms. The integration of cultivation results, natural abundance data from inoculum communities, and DTE modeling represents an important step forward in quantifying the risk associated with DTE efforts to isolate valuable taxa from new sources, or repeating isolation from the same locations. We hope variations of this approach will be incorporated into wider community efforts to invest in culturing the uncultured.

## Acknowledgments

We would like to thank Dr. Nancy Rabalais for her comments and edits on a very early draft of this manuscript. Portions of this research were conducted with high-performance computing resources provided by Louisiana State University (http://www.hpc.lsu.edu) and the University of Southern California (http://hpc.usc.edu).

## Funding Information

This work was supported by the Department of Biological Sciences at Louisiana State University, a Louisiana Board of Regents grant to JCT (LEQSF(2014-17)-RD-A-06), a National Academies of Science, Engineering, and Medicine Gulf Research Program Early Career Fellowship to JCT, and the Dornsife College of Letters, Arts and Sciences at the University of Southern California. BT was partially funded by NERC award NE/R010935/1 and by the Simons Foundation BIOS-SCOPE program. The funders had no role in study design, data collection, and interpretation, or the decision to submit the work for publication.

## Author Contributions

MWH led sample collections, cultivation experiments, nucleic acid extraction, amplicon sequencing, and analyses; VCL, DMP, JLW, and AML supported sample collections, cultivation experiments, and nucleic acid extractions; MWH and JCT conducted cultivation comparisons and phylogenetic analyses; BT developed the viability model and led the statistical analyses; CC derived the cell-specific growth rate equations incorporating viability; JCT designed the study and assisted with sample collections and model refinement; MWH, BT, and JCT led manuscript writing; and all authors contributed to and reviewed the text.

## Notes

https://figshare.com/account/home#/projects/78987

https://github.com/thrash-lab

### Competing Interest Statement

The authors have declared no competing interest.

### Summary of Updates

Changes made in response to reviewer comments.

