## Supplemental Information for "Expanding the diversity of bacterioplankton isolates and modeling isolation efficacy with large scale dilution-to-extinction cultivation"

**Supplemental Text**
Math for back calculating the per-cell division rates in Henson et al. 2018 assuming 15%--55% viability.

Denote the initial cell count as $X_0$, and the cell count in $n^{th}$generation as$X_n$. If all the cells are viable, per each generation, all the cells are doubling:

$$X_n = X_{n-1} \cdot 2 \quad \text{(eq. S1.1)}$$

And therefore, after $n$generations, the cell count $X_n$would be:

$$X_n = X_0 \cdot 2^n \quad \text{(eq. S1.2)}$$

When not all the cells are viable, for each generation, instead of simply doubling the whole population, only a portion of the cells (denoted as$V < 1$) replicates. In this case:

$$X_n = X_{n-1} \cdot (1 + V) \quad \text{(eq. S1.3)}$$

Here we can see that (eq. 3.1) is actually a special case of (eq. 3.3), when $V = 1$,

$$X_n = X_{n-1} \cdot (1 + 1) = X_{n-1} \cdot 2$$

When $V < 1$, after $n$ generations:

$$X_n = X_0 \cdot (1 + V)^n \quad \text{(eq. S1.4)}$$

Denoting the total time to get $n$ generations as $t$, which means the number of generations per unit time, also known as the per-cell division rate, is $\mu = n/t$. Rewriting (eq. 3.4), we have:

$$X_n = X_0 \cdot (1 + V)^{\mu \cdot t} = X_0 \cdot 2^{\mu \cdot log_2(1+V) \cdot t} \quad \text{(eq. S1.5)}$$

According to (eq. 3.5), a cell population of $V$viability and $\mu$ per cell division rate, the population division rate is:

$$\mu_{population} = \mu \cdot log_2(1 + V) \quad \text{(eq. S1.6)}$$

Henson et al. 2018 show that the optimal population division rate of SAR11 LD12 is $\mu_{population} = 0.5\ day^{-1}$, assuming the viability of LD12 is $V = 0.15$, by plugging in these two numbers to (eq. S1.6), we can get the per-cell division rate of LD12 is:

$$\begin{aligned} \mu &= \mu_{population}/log_2(1 + V) \\ &= 0.5\ day^{-1}/log_2(1 + 0.15) \\ &= 0.5\ day^{-1}/0.20163 \\ &= 2.48\ day^{-1} \end{aligned}$$

If assuming $V = 0.55$, $\mu = 0.79\ day^{-1}$. Therefore, given the range of the viability of LD12 is between 15%--55%, the corresponding per cell division rate is between 0.79 divisions to 2.48 divisions per day.

**Supplemental Table**
Supplemental Table S1 is a spreadsheet, Table_S1.xlsx. This includes MWH media recipes, ASV and OTU tables, taxonomic and relative abundance information for all isolates, NMDS information, biogeochemical data for the samples, and the BLAST results of isolate hits. Available at https://doi.org/10.6084/m9.figshare.12142113.

**Supplemental Figures follow, with captions embedded**

**Figure S1**

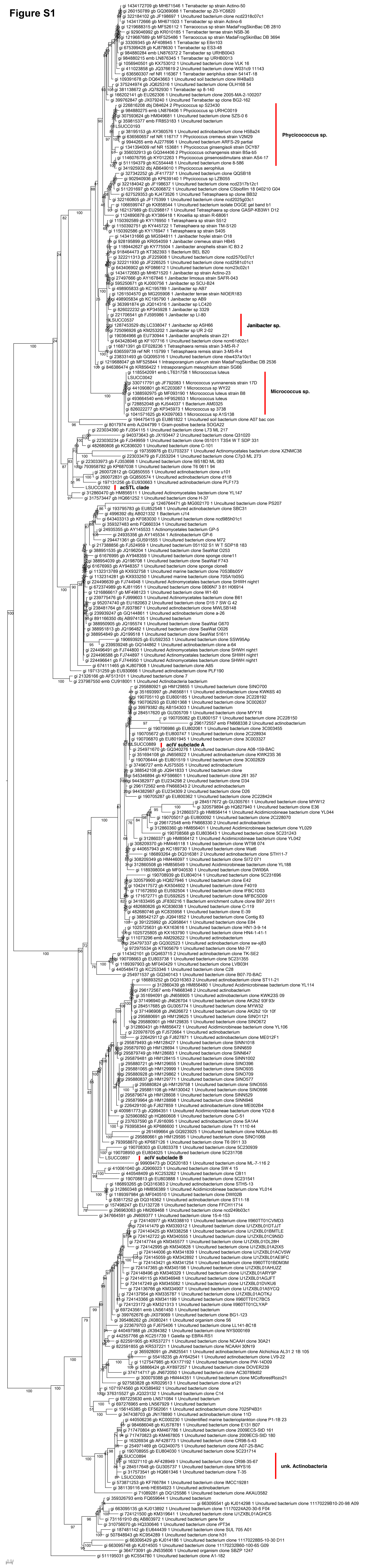
Phycicoccus sp.
Janibacter sp.
Micrococcus sp.
acSTL clade
acIV subclade A
acIV subclade B
unk. Actinobacteria

### Figure S2

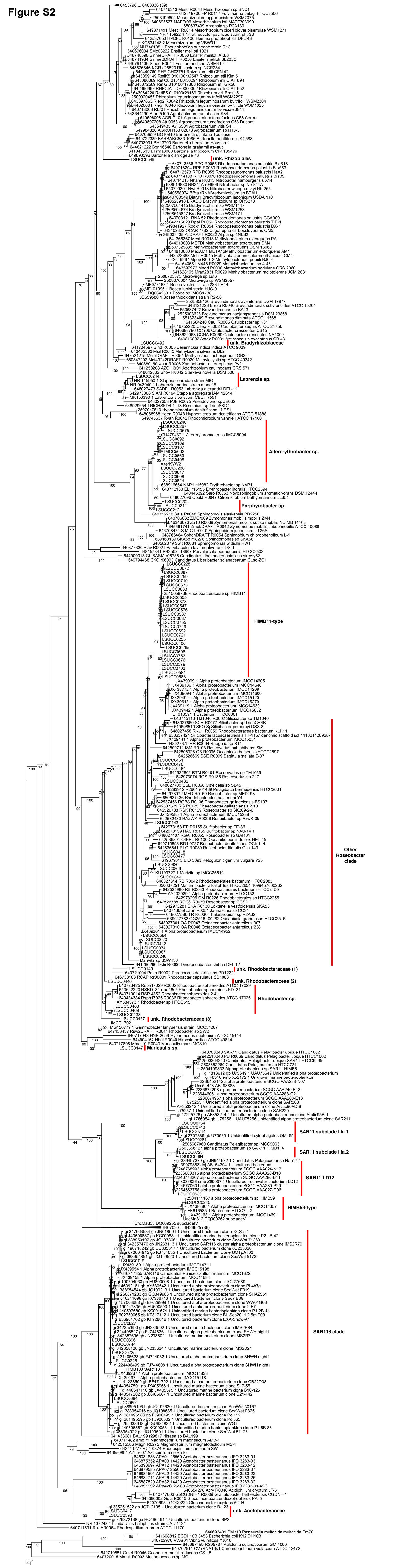
unk. Rhizobiales
unk. Bradyrhizobiaceae
Labrenzia sp.
Altererythrobacter sp.
Porphyrobacter sp.
HIMB11-type
Other Roseobacter clade
unk. Rhodobacteraceae (1)
unk. Rhodobacteraceae (2)
Rhodobacter sp.
unk. Rhodobacteraceae (3)
Maricaulis sp.
SAR11 subclade IIIa.1
SAR11 subclade IIIa.2
SAR11 LD12
HIMB59-type
SAR116 clade
unk. Acetobacteraceae

### Figure S3

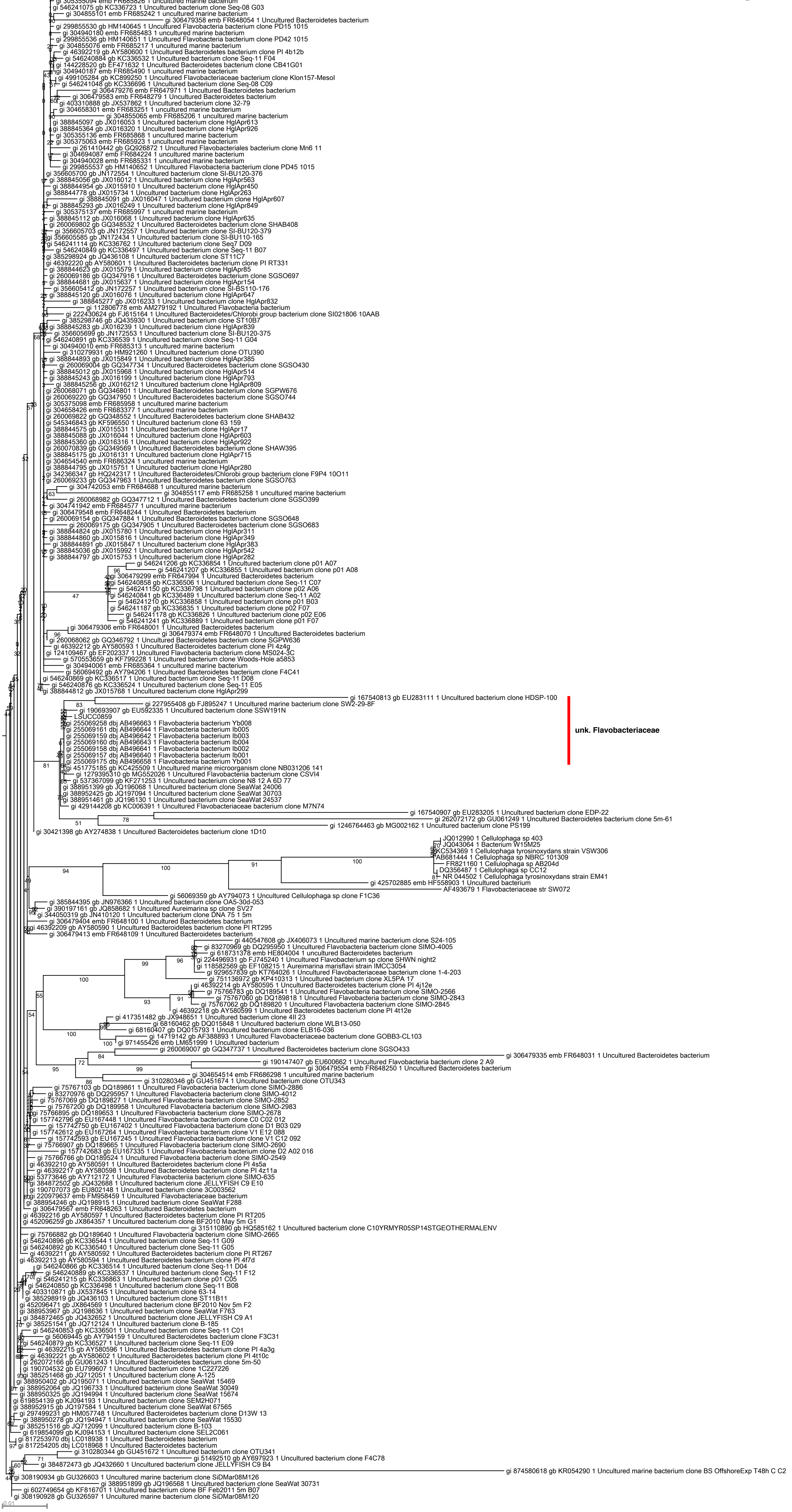

### Figure S4

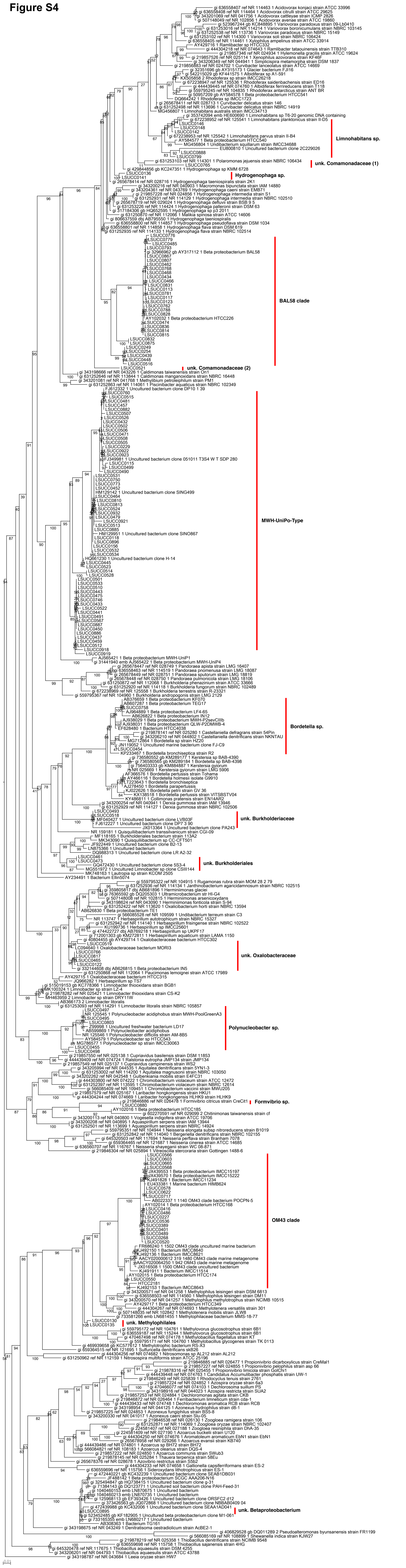

Limnohabitans sp.
unk. Comamonadaceae (1)
Hydrogenophaga sp.
BAL58 clade
unk. Comamonadaceae (2)
MWH-UniPo-Type
Bordetella sp.
unk. Burkholderiaceae
unk. Burkholderiales
unk. Oxalobacteraceae
Polynucleobacter sp.
Formivibrio sp.
OM43 clade
unk. Methylophilales
unk. Betaproteobacterium

**Figure S5**

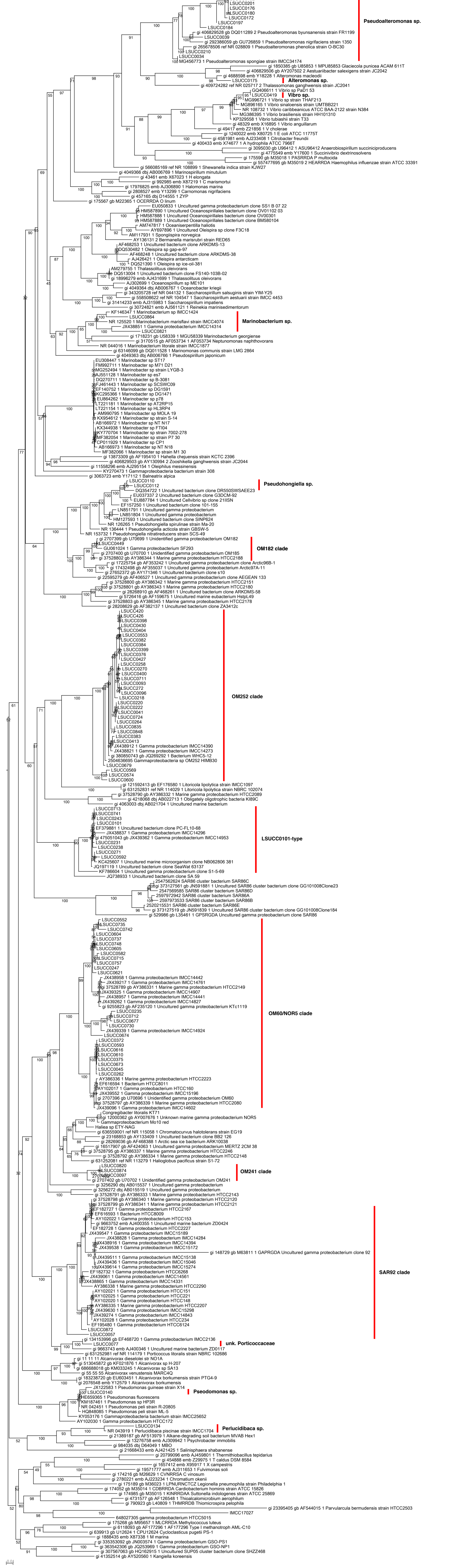

Pseudoalteromonas sp.
Alteromonas sp.
Vibro sp.
Marinobacterium sp.
Pseudohongiella sp.
OM182 clade
OM252 clade
LSUCC0101-type
OM60/NOR5 clade
OM241 clade
SAR92 clade
unk. Porticoccaceae
Pseodomonas sp.
Perlucidibaca sp.

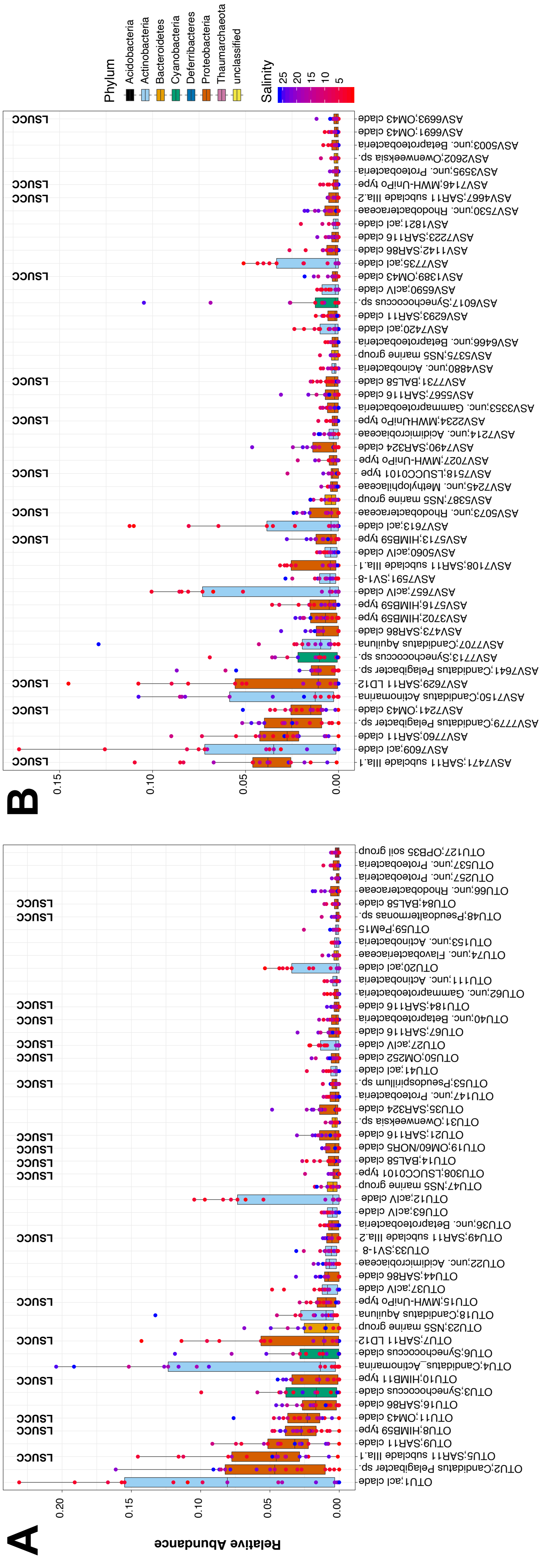


**Figure S6.** Rank abundance plot of the top 50 most abundant (A) OTUs and (B) ASVs across all sites. The boxes indicate the interquartile range (IQR) of the data, with vertical lines indicating the upper and lower extremes according to 1.5 x IQR. Horizontal lines within each box indicate the median. The underlying data points are each individual OTU's sample relative abundances. The shade of the dot represents the salinity, while the color of the box is the Phylum of the OTU. OTUs with cultured representatives from the LSU culture collection are labeled with one LSUCC isolate.

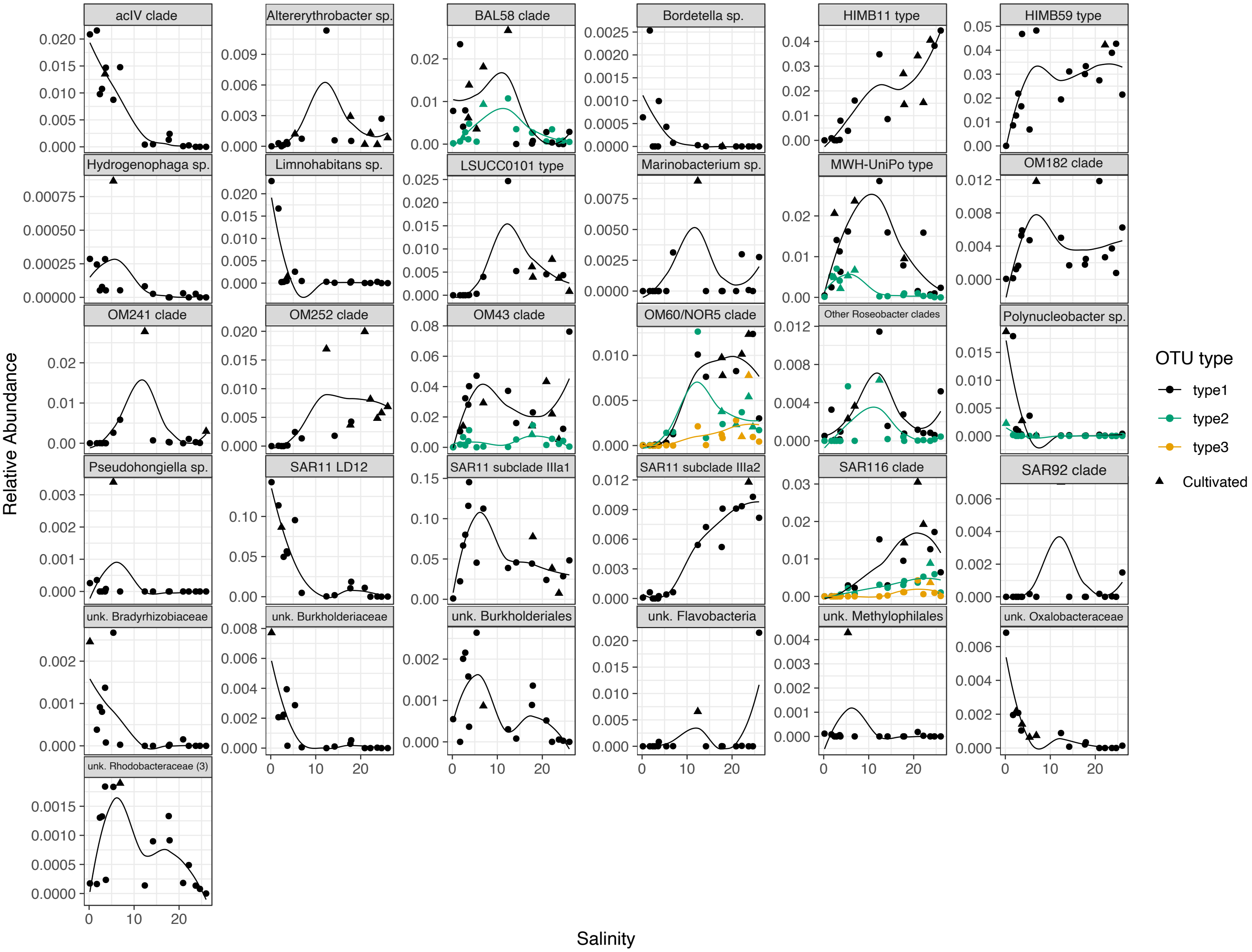


**Figure S7.** Relative abundance of cultivated OTUs according to site salinity. The color of the line represents the different OTUs classified within the LSUCC group. Non-linear regressions have been added for reference.

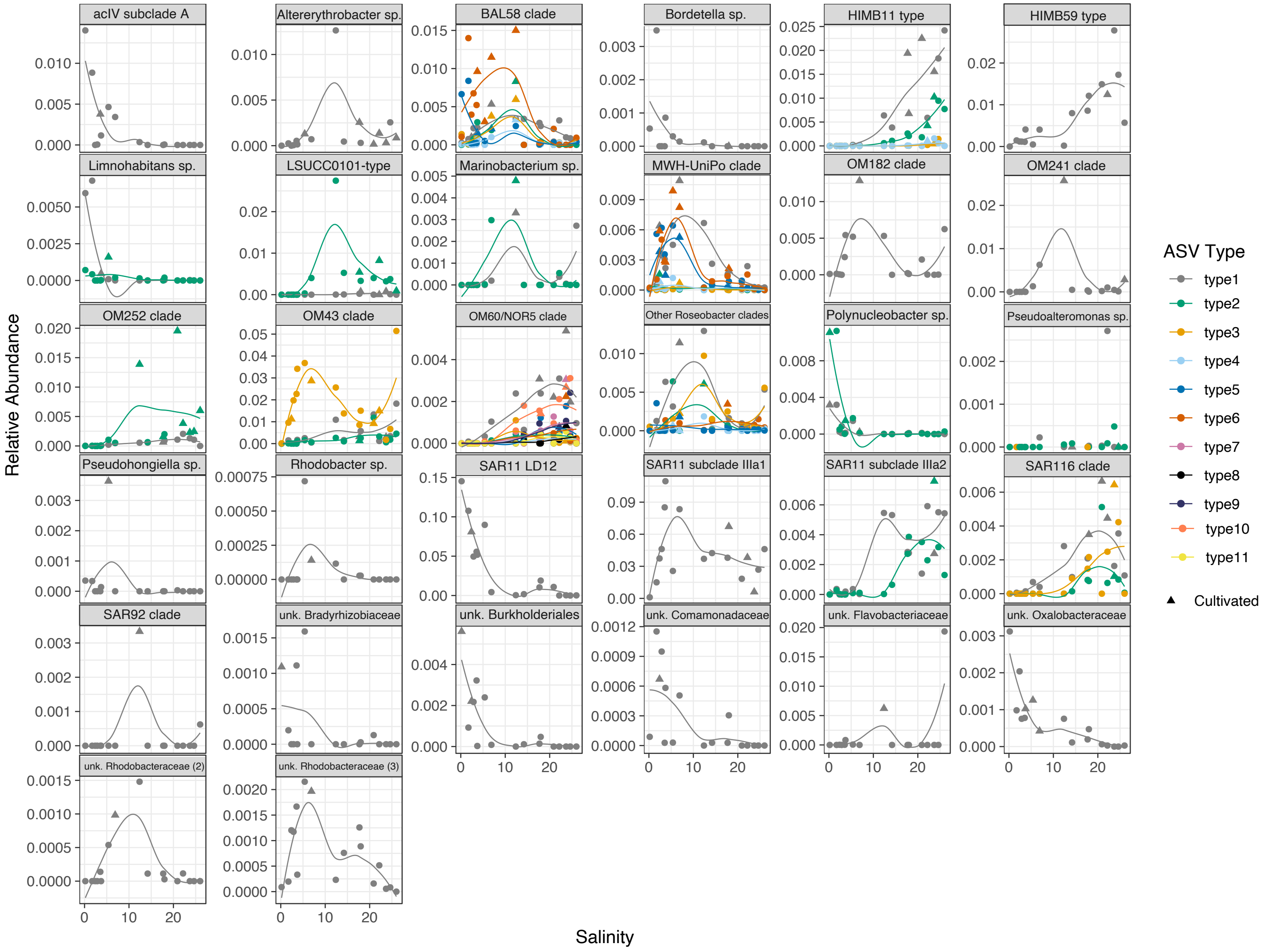


**Figure S8.** Relative abundance of ASVs with a cultured LSUCC representative according to site salinity. The color of the line represents the different ASVs classified within the LSUCC group. Non-linear regressions have been added for reference. Arrows represent sites where isolates were cultivated.

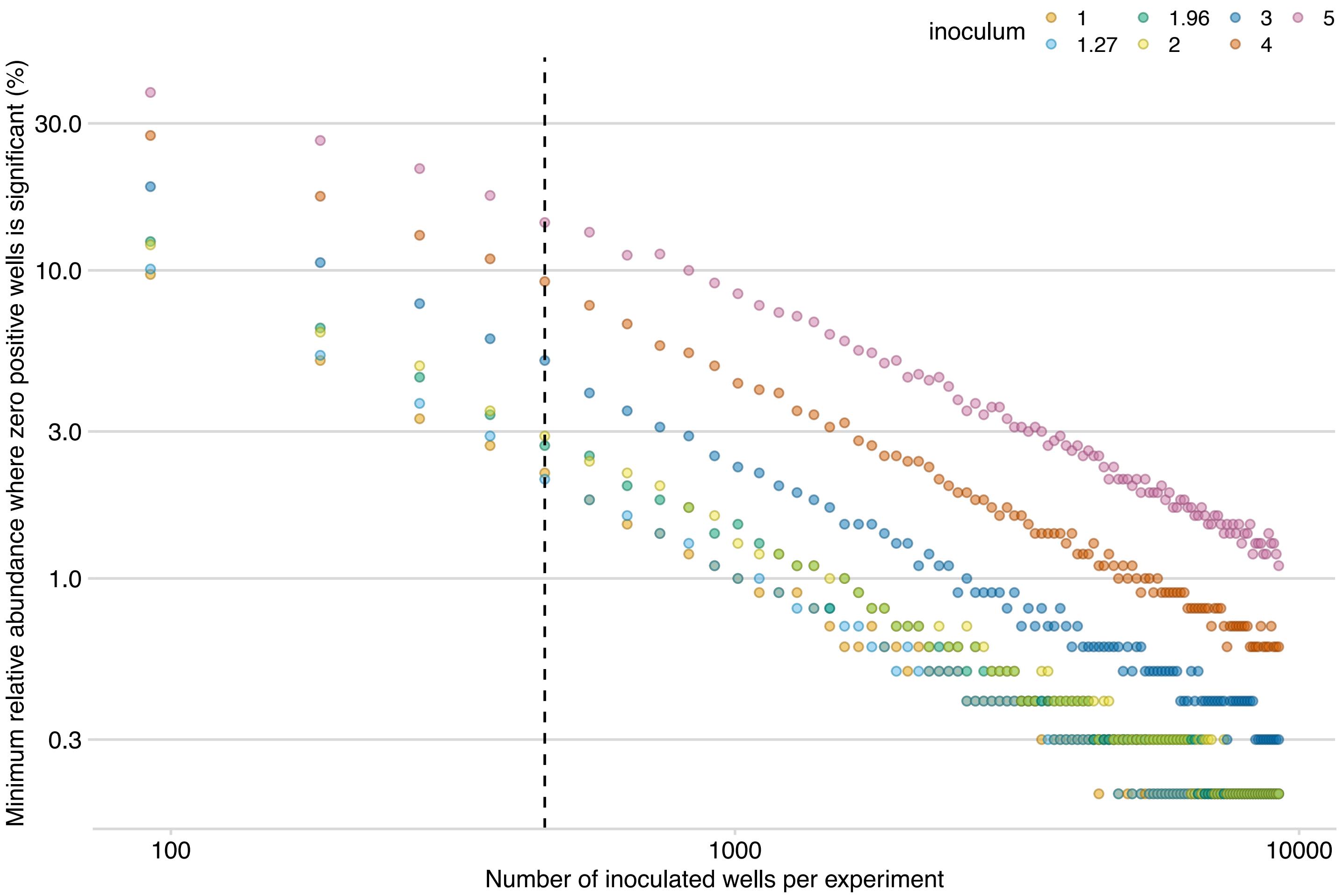


**Figure S9.** Sensitivity testing of the viability model to determine minimum numbers of wells required to overcome false negatives associated with differing levels of taxon relative abundance. Different inoculum densities (λ) are depicted with colors. The dotted line indicates 460 wells which corresponded to the experiments in this study.

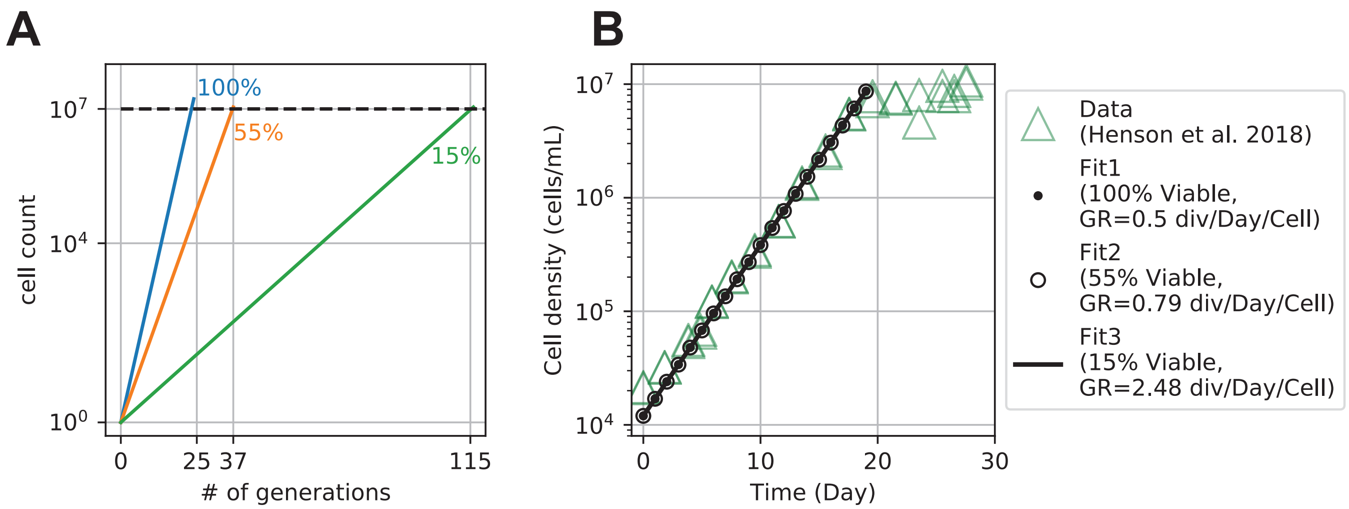


**Figure S10.** Logarithmic growth with different percentages of cell viability. Under different cell viabilities, such as 100%, 55% and 15% shown in (A), the culture can have logarithmic growth in the population size if we assume no cell death. It takes around 25 generations to have 10^7fold expansion of population size when all cells are active (100% viable), whereas when 15% of the population is actively duplicating, it takes around 115 generations to reach the same population size expansion. B) For the SAR11 LD12 strain LSUCC0530, the previously measured population doubling rate was 0.5/day (29). If all the cells are active (100% viability), each cell within the population will double 0.5 times per day (Fit 1 in B). However, with 15% viability, the actual active cell doubling rate is 2.48 doublings day-1 (Fit 3 in B).
